# Beyond miRNA cargo profiles: anti-inflammatory roles of extracellular vesicle-enriched miRNAs derived from human intervertebral disc cells unveiled by functional testing

**DOI:** 10.1101/2025.10.08.681183

**Authors:** Li Li, Aiwei Sun, Saber Ghazizadeh, Hadil Al-Jallad, Kirby Upshaw, Saleh Alfaisali, Jean Ouellet, Peter Jarzem, Hosni Cherif, Lisbet Haglund

## Abstract

Intervertebral disc (IVD) degeneration is a leading cause of chronic low back pain and a major contributor to global disability. Understanding the molecular mechanisms underlying this condition is essential for developing targeted therapies. Among these mechanisms, microRNAs (miRNAs) have emerged as critical post-transcriptional regulators of gene expression in IVD cells, influencing key processes such as extracellular matrix (ECM) homeostasis, inflammatory signalling, and cellular senescence. Extracellular vesicles (EVs), which transport miRNAs between cells, represent a promising avenue for therapeutic intervention. However, the composition of their miRNA cargo across different stages of disc degeneration remains inadequately characterized. We isolated EVs from primary human IVD cells derived from non-degenerate, mildly-degenerate, and severely degenerate tissues, and performed small RNA sequencing to profile their miRNA content. Bioinformatic analyses revealed enrichment in pathways related to ECM-receptor interaction, focal adhesion, inflammation, and cell cycle regulation. Notably, let-7b-5p and miR-100-5p were among the most abundant miRNAs and were significantly lower in EVs from degenerate discs. Functional assays demonstrated that transfection of IVD cells with let-7b-5p or miR-100-5p mimics individually suppressed IL-1ß expression at both mRNA and protein levels, confirming their anti-inflammatory roles. Strikingly, co-delivery of both miRNAs enhanced suppression of pro-inflammatory mediators, reduced senescence-associated *p16* expression, and upregulated *TIE2* mRNA, indicating synergistic effects in promoting a regenerative cell phenotype. These findings highlight the regulatory roles of EV-enriched let-7b-5p and miR-100-5p in modulating inflammation and senescence in IVD cells, and underscore the potential of miRNA-loaded EVs as cell-free regenerative therapies for disc degeneration.

## 1 Introduction

Intervertebral disc (IVD) degeneration is a major contributor to chronic low back pain (LBP), the leading cause of disability worldwide, particularly affecting aging populations in developed countries.^1^ In addition to its impact on quality of life, IVD degeneration imposes a substantial economic burden through increased healthcare demands and reduced workforce productivity.^1,2^ Epidemiological studies and magnetic resonance imaging analyses have underscored the progressive nature of this condition, highlighting the urgent need for innovative therapeutic strategies.^3,4^

Extracellular vesicles (EVs) have emerged as promising cell-free tools in regenerative medicine, especially for degenerative diseases such as IVD degeneration.^5-7^ EVs are lipid bilayer-bound particles secreted by cells that carry bioactive cargo, including microRNAs (miRNAs), proteins, and lipids.^8^ Their ability to mimic the functional properties of their parent cells makes them ideal candidates for delivering targeted molecular therapies to damaged tissues.^6,9^ Among these cargo molecules, miRNAs are key regulators of gene expression and reflect the homeostatic state of tissues.^10-12^

Recent studies have shown that EVs derived from human mesenchymal stem cells (MSCs), enriched with anti-inflammatory miRNAs, can suppress pro-inflammatory cytokine expression in IVD cells, potentially slowing degeneration.^5^ Specific miRNAs such as miR-155-5p and miR-27a have been implicated in activating inflammatory pathways, while their inhibition leads to reduced cytokine levels.^13,14^ Other miRNAs, including miR-149-5p and miR-99a-3p, have been shown to inhibit matrix metalloproteinases and prevent extracellular matrix (ECM) degradation.^15^ Furthermore, combinations of miRNAs like miR-181a-5p, miR-92a-3p, miR-21-5p, and miR-186-5p delivered via EVs have demonstrated the ability to reduce senescence markers and inflammatory mediators in both human cell cultures and animal models.^16^ These findings suggest that profiling miRNAs associated with tissue health could help identify therapeutic candidates capable of mitigating degeneration, chronic pain, and functional impairment.^17,18^ Engineered EVs carrying specific miRNAs offer a controlled approach to modulate cellular responses and promote tissue regeneration in the IVD microenvironment.^19-21^

Despite these promising developments, significant gaps remain in our understanding of the miRNA cargo within EVs derived directly from human IVD cells. Most existing research has focused on MSC-derived EVs, with limited investigation into the miRNA profiles of EVs from IVD cells themselves.^7,22^ Given the role of EV-associated miRNAs in regulating biological processes relevant to IVD degeneration, a comprehensive profiling of these molecules is essential. To date, no study has systematically examined the miRNA content of EVs from human IVD cells isolated from tissues at varying stages of degeneration using standardized culturing, purification, sequencing, and bioinformatic analysis protocols. Additionally, functional studies evaluating the roles of these miRNAs in IVD health are sparse, and no prior work has employed a function prediction-guided screening strategy to assess their therapeutic potential.

Therefore, the objectives of this study were to: (1) characterize the miRNA profiles of EVs derived from human IVD cells across different grades of tissue degeneration, and (2) evaluate the therapeutic potential of key miRNAs using a targeted, function prediction-guided screening approach.

## 2 Materials and Methods

### 2.1 Study workflow

Total RNA was extracted from purified EVs, and small RNA was purified from the bulk RNA. Small RNA sequencing was conducted after the quality test. Bioinformatics was used to predict the function of detected miRNAs. Five miRNAs of interest were selected for further functional assessments. (**Figure 1A**) The functional assessments comprised the evaluation of the regulatory roles and the expression of selected miRNAs at the gene and protein levels. (**Figure 1B**) The expression of nucleus pulposus (NP) markers, inflammatory mediators, and senescence markers were evaluated to determine the effect on gene expression level. (**Figure 1B(a)**) The production of inflammatory mediators was evaluated to validate the anti-inflammatory effects of selected miRNAs at the protein level. (**Figure 1B(b)**)

**Figure 1.**
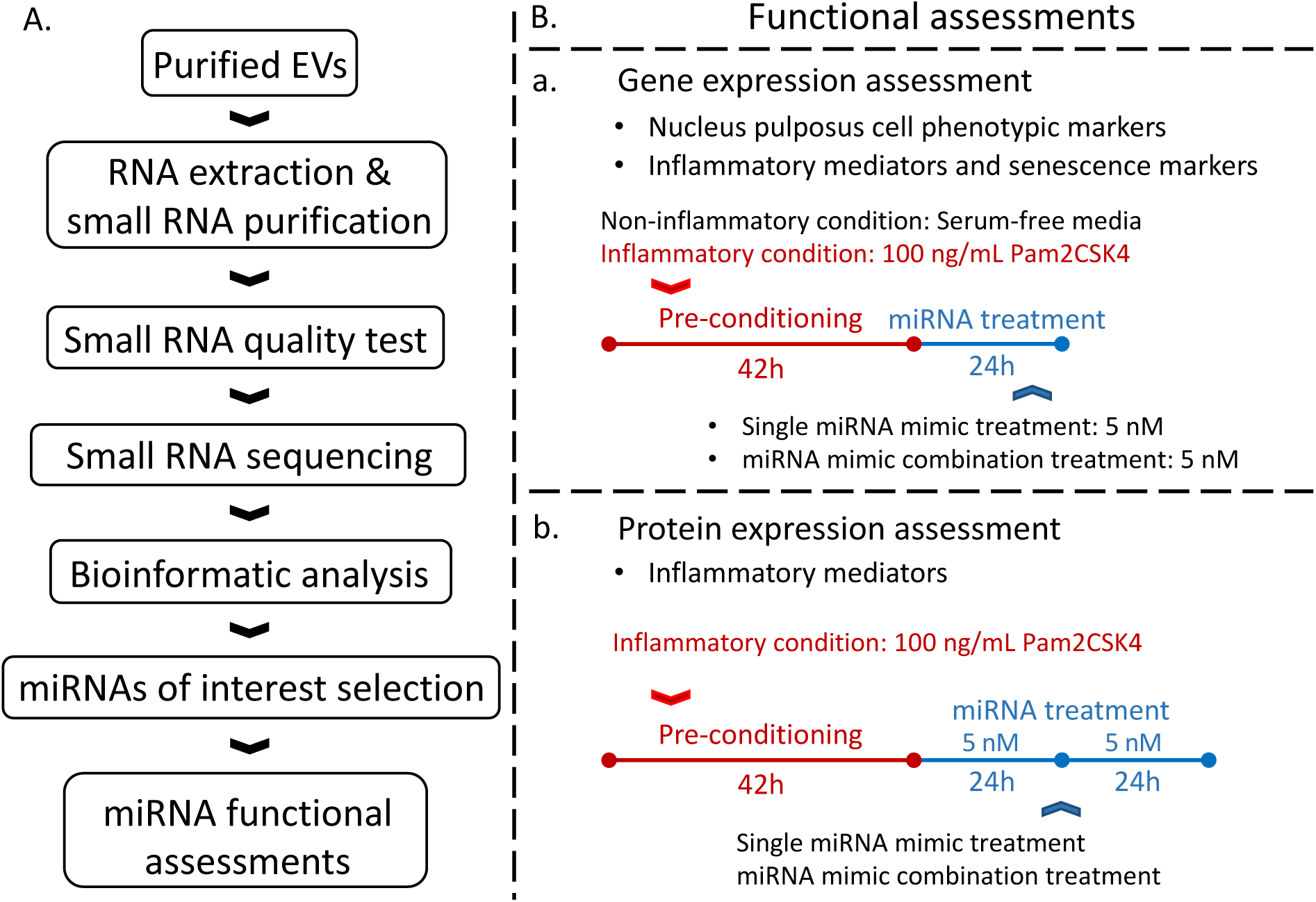
Schematic overview of the study workflow. **A**. Study workflow includes RNA extraction and small RNA purification, small RNA quality test, small RNA sequencing, bioinformatic analysis, selecting the miRNAs of interest, and functional assessments. **B.** Functional assessments including (**a**) gene expression of NP cell phenotypic markers, inflammatory mediators, and senescence markers. (**b**) protein expression of inflammatory mediators.

### 2.2 Sample preparation

This study was conducted in strict compliance with ethical protocols approved by McGill University’s Institutional Review Board (IRB #00010120). Purified EVs used for small RNA sequencing were obtained as previously described.^23^ Samples from additional donors were included in this study for the miRNA functional assessments. Donor and patient demographic information are presented in **Table 1**.

**Table 1.**
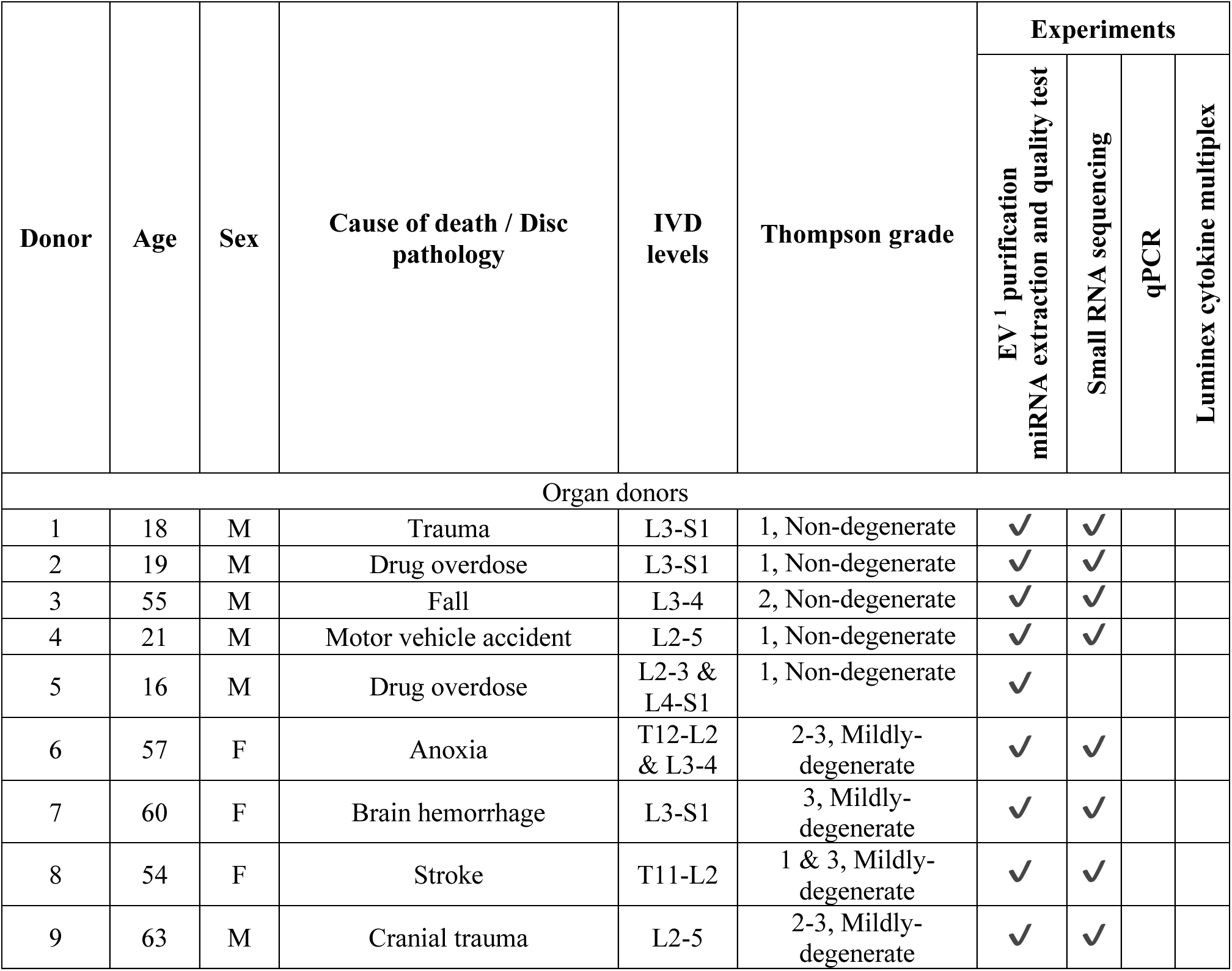
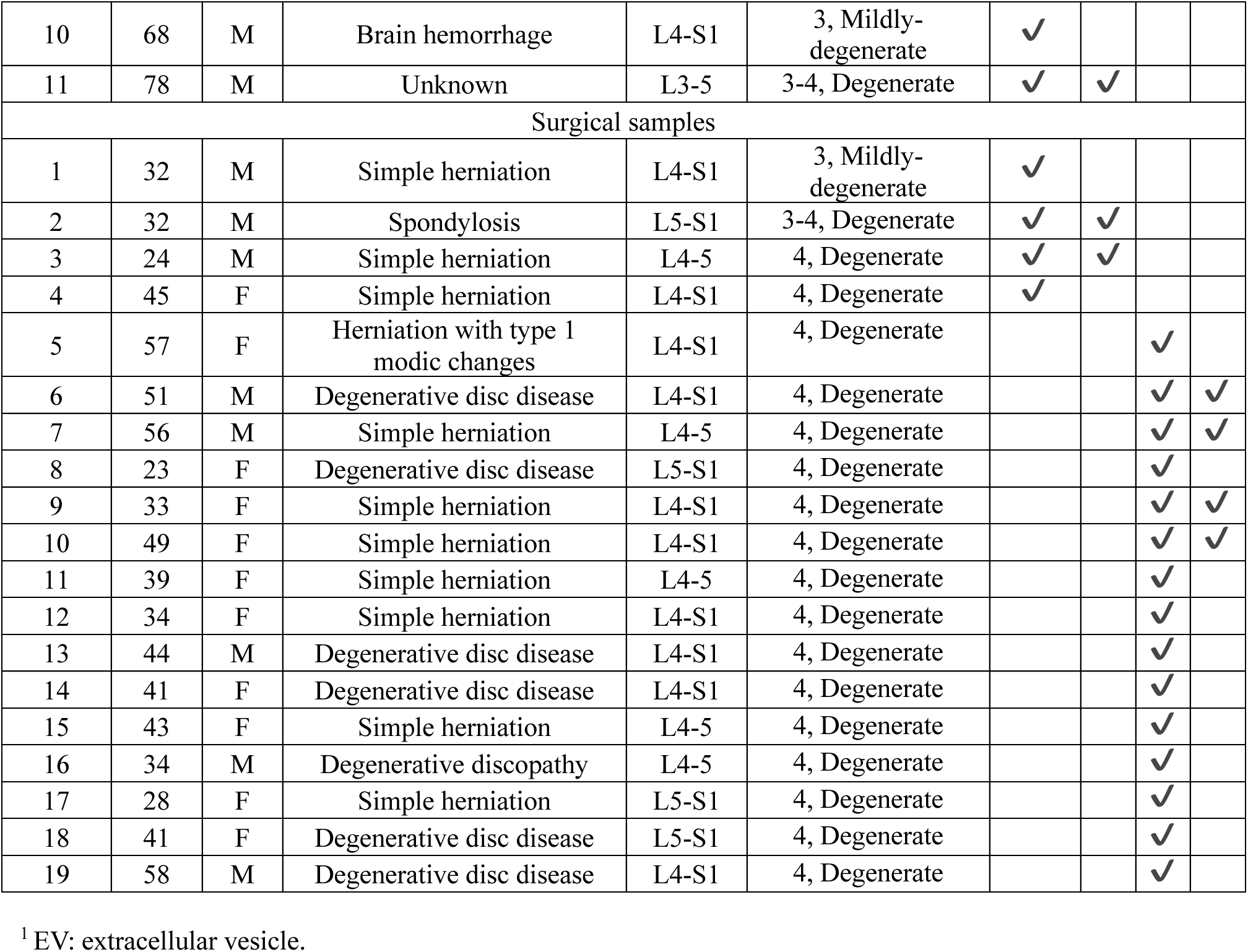
Donor and patient information and applications.

#### MiRNA purification

EV samples (approximately 1.84 × 10^10^ particles/mL, 50 µL) from 15 donors (4-6 biological replicates) were used. EVs were purified using size-exclusion chromatography purification in phosphate-buffered saline (PBS) and fractions 7-9 were pooled. RNA was extracted and miRNA was purified from the EV samples with a miRNeasy micro kit (Cat. #217084, QIAGEN, Toronto, ON, Canada) following the manufacturer’s instructions. MiRNAs were stored at -80°C for further analysis.

#### MiRNA quality test

The quantity and quality of miRNAs were evaluated with a Qubit^TM^ RNA High Sensitivity Assay Kit (ThermoFisher Scientific, Saint-Laurent, QC, CA) and the Agilent automated electrophoresis portfolio, including the Agilent 2100 Bioanalyzer system and Small RNA Analysis kit with RNA 6000 Pico Chips. (Agilent Technologies, Mississauga, ON, Canada). All miRNA samples passed quality control.

### 2.3 Small RNA sequencing and data processing

#### Library preparation and quality test

Small RNA sequencing libraries were prepared using QIAseq miRNA Library Kit (12-nt unique molecular identifiers (UMIs)) (REF. #1103677, LOT. #76602338, QIAGEN, Toronto, ON, Canada) following the manufacturer’s instructions. Sequencing was performed on a NextSeq 500 system (Illumina, San Diego, CA, USA), obtaining around 10 million reads per sample. Libraries were quantified using a Qubit^TM^ dsDNA Quantification High Sensitivity Assay Kit (ThermoFisher Scientific, Toronto, ON, Canada) and assessed by Agilent 2100 Bioanalyzer using an Agilent High Sensitivity DNA Kit (Agilent Technologies, Mississauga, ON, Canada). All libraries passed the quality test.

#### Data processing

Raw FASTQ files were analyzed using QIAGEN’s own analytical pipeline RNA-seq Analysis Portal 5.1 (https://geneglobe.qiagen.com/ca/analyze) using the provided *Homo sapiens* (GRCh38.103) genome and miRBase v22.0 database. Samples passed quality control if > 1% of UMI reads were annotated to miRBase v22.0 records. Eleven out of 15 samples (3-4 donors per group) passed the quality control. Raw data of miRNA expression in count per million (CPM) of the 11 samples was exported for further analysis.

### 2.4 Pathway and annotation analysis for miRNA function prediction

#### Function prediction

The enriched pathway and annotation analyses were performed by using *DIANA-miRPath v4.0*, a web server that offers numerous analysis options, including Kyoto Encyclopedia of Genes and Genomes (KEGG) and Gene Ontology (GO) analyses, for the target-based functional analysis of miRNAs.^24,25^ We performed miRNA-centric analysis for KEGG and GO pathways and annotations, and included only experimentally supported human miRNA targets on protein-coding transcripts by using the TarBase v8.0 database to avoid false-positive predictions. We applied direct TarBase targets without using a secondary target annotation source when no significant interactions were found for certain miRNAs and did not include long non-coding RNA targets. The miRNA annotation (or miR-ID) was based on miRBase v22.1 database. We used the Pathways Union merging method combined with the Classic Analysis testing method to assess the combined action of selected miRNAs. False discovery rate (FDR) correction and a *p*-value threshold of ≤ 0.05 were applied. Identical or duplicate reads in low-abundance miRNAs are a common observation in small RNA sequencing, particularly when miRNA concentrations are low. This phenomenon can arise from technical biases, such as those introduced during the PCR amplification step.^26^ We observed increased identical reads in miRNAs with CPM ≤ 300. To minimize false positive results, we focused the functional prediction on miRNAs with CPM > 300. *Evaluation metrics* and *algorithmic process* are described in the *Supplementary Methods*.

### 2.5 Cell culture and induction of an inflammatory microenvironment

#### Cell culture

Human IVD cell isolation was performed as previously published.^23,27^. We used cells from surgical samples for functional testing, which were obtained from a mixture of NP and inner annulus fibrosus (AF) tissue. Isolated cells were seeded at one million cells per T75 cell culture flask (Sarstedt, TC Flask T75, Stand, Vent. Cap, Germany). Cells were cultured in Dulbecco’s Modified Eagle Medium (DMEM) (Sigma-Aldrich, Oakville, ON, Canada) containing 2.25 g/L glucose. The glucose level was selected for consistency with the EV collection media.^23^ The media was further supplemented with 10% (v/v) fetal bovine serum (FBS), 50 µg/mL Gentamicin, 2 mM GlutaMAX^TM^, and 50 mM HEPES (ThermoFisher Scientific, Toronto, ON, Canada). The cultures were maintained at 37 °C in a 5% CO_2_ environment for 1-2 weeks. The culture media was refreshed twice per week until cells reached approximately 80% confluence in passage 1. The cells then passaged and reseeded into a 6-well-plate (Sarstedt, Germany) at a density of 240,000 cells/well with 3 mL/well media and maintained in the same condition overnight. To minimize the effects of isolation and expansion in monolayer cultures, we only used cells at passage 1 for expansion and passage 2 for functional testing in this study.

#### Toll-like receptor (TLR) activation

Synthetic diacylated lipopeptide Pam2CSK4 (Pam2) (Cat. #tlrl-pm2s-1, InvivoGen, San Diego, CA, USA), a potent activator of TLR2 and the pro-inflammatory transcription factor NF-κB, was used to induce an inflammatory microenvironment.^28-30^ Monolayer human IVD cells were rinsed with 1 mL/well sterile PBS three times. Pam2 (100 ng/mL) was applied in 3 mL/well serum-free media (DMEM (Sigma-Aldrich, Oakville, ON, Canada) containing 2.25 g/L glucose, supplemented with 50 µg/mL Gentamicin, 2 mM GlutaMAX^TM^, and 50 mM HEPES (ThermoFisher Scientific, Toronto, ON, Canada)) and incubated for 42 h at 37 °C in a 5% CO_2_ environment.^29^

### 2.6 Gene expression assessment

#### MiRNA transfection

Human miRCURY LNA miRNA Mimics (QIAGEN, Toronto, ON, Canada), let-7a-5p, let-7b-5p, let-7f-5p, miR-16-5p, and miR-100-5p, were prepared following the manufacturer’s instructions. Single-miRNA treatment (5 miRNA mimics, respectively) and miRNA combination treatment (the combinations of two miRNAs of let-7b-5p, miR-16-5p, miR-100-5p, and the combinations of all of them, respectively) were prepared at a working concentration of 5 nM for each miRNA mimic^31^ in the solution using the above-mentioned serum-free complete media. Human IVD cells were rinsed with 1 mL/well sterile PBS for 3 times and transfected with 3 mL/well miRNA mimic solution for 24 h using 4 µL/mL HiPerFect transfection reagent (QIAGEN, Toronto, ON, Canada) according to the manufacturer’s protocols. (**Figure 1B(a)**)

#### Quantitative polymerase chain reaction (qPCR)

IVD cells were lysed in TRIzol^TM^ reagent (ThermoFisher Scientific, Toronto, ON, Canada) and RNA extraction was performed following the manufacturer’s instructions. Reverse transcription and qPCR procedures were conducted as previously described.^32-34^ Briefly, complementary DNA (cDNA) synthesis was conducted with a High-Capacity cDNA Reverse Transcription Kit (ThermoFisher Scientific, Toronto, ON, Canada) using Applied Biosystems^®^ Veriti^®^ 96-Well Fast Thermal Cycler (ThermoFisher Scientific, Toronto, ON, Canada). QPCR was performed in PROGENE^®^ 96-Well Half-Skirt ABI Fast PCR Plates (UltiDent Scientific, Montreal, QC, Canada) with PROGENE^®^ Adhesive Routine PCR Sealing Film (UltiDent Scientific, Montreal, QC, Canada) using an Applied Biosystems^®^ StepOnePlus^TM^ Real-Time PCR System (ThermoFisher Scientific, Toronto, ON, Canada) with PowerUp^TM^ SYBR^TM^ Green Master Mix (ThermoFisher Scientific, Toronto, ON, Canada). Primers are presented in **Table 2**. The melting curves were examined to exclude the risks of false amplifications. Data were normalized to the geometric mean of the reference genes (*beta-ACTIN* and *28S rRNA*)^35^ and presented in fold-changes calculated with the 2^-ΔΔCt^ method.^36^

**Table 2.**
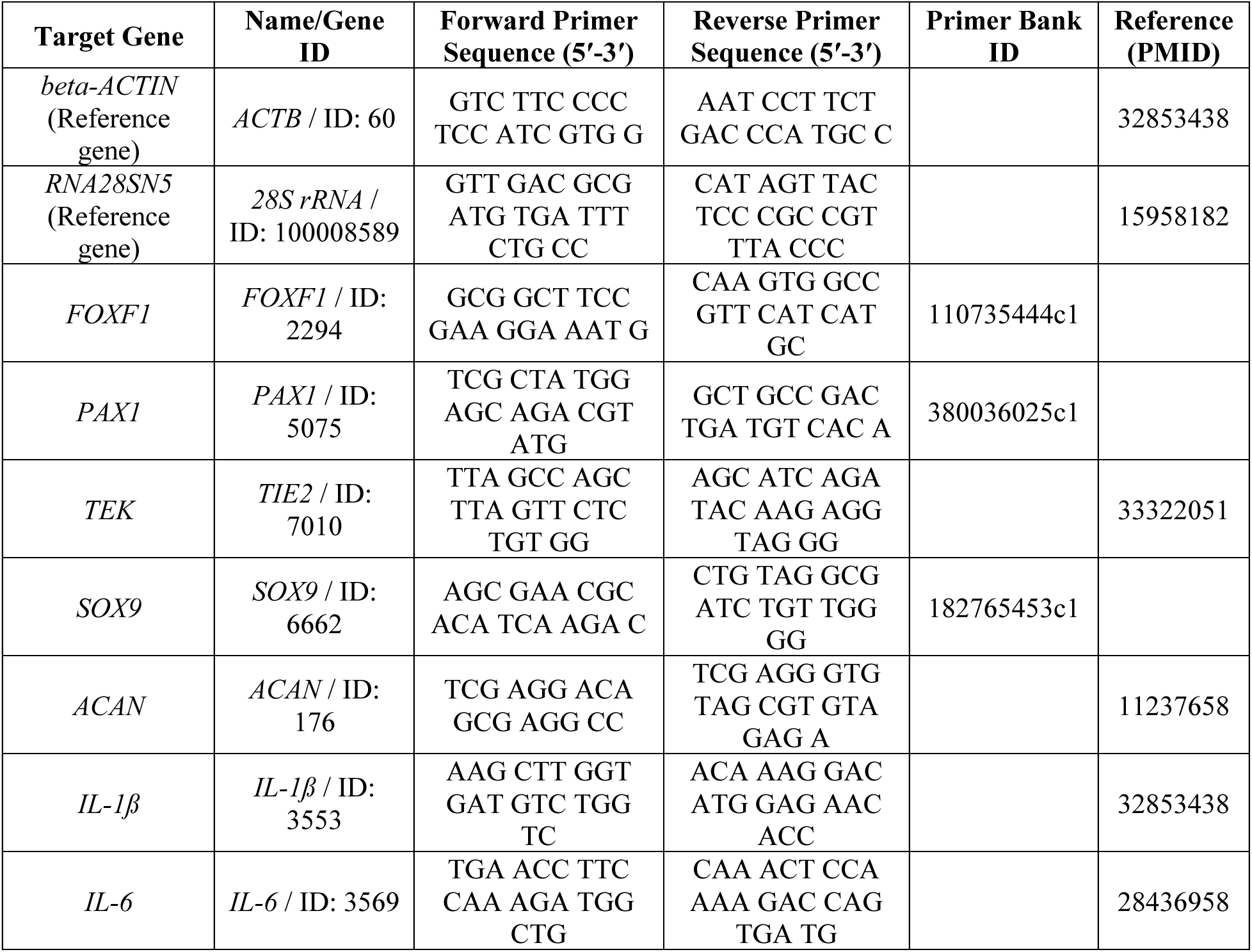
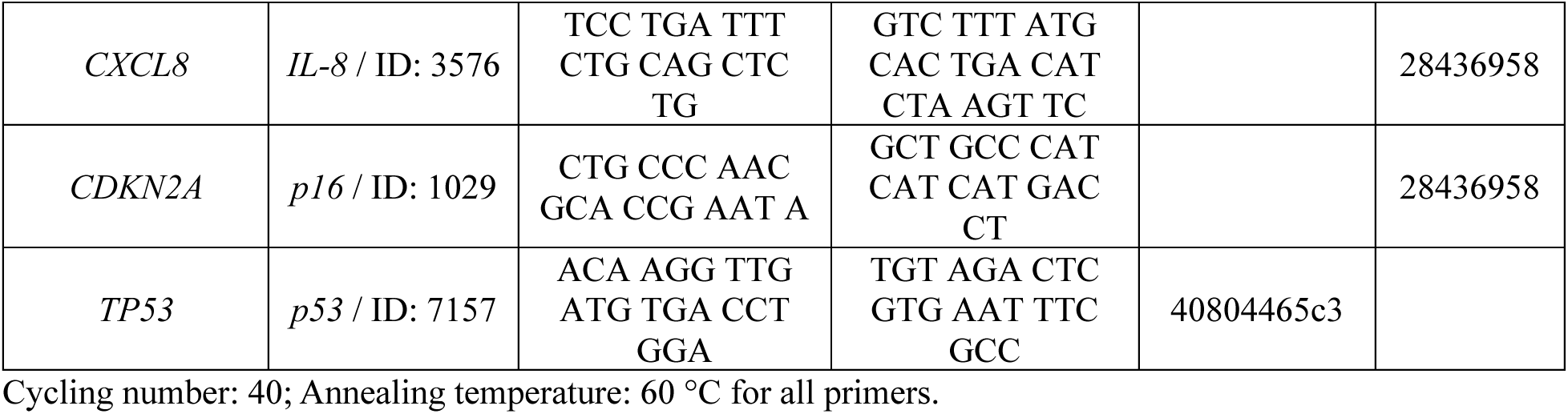
qPCR primer list.

In addition, our recent single-cell RNA-sequencing study has shown that NP and inner AF cells, both of which exhibit a chondrocyte-like phenotype, share similar transcriptional profiles.^37^ Consequently, commonly used NP phenotypic markers, such as *TEK receptor tyrosine kinase (TIE2)* and *Aggrecan (ACAN)*, are also robustly expressed in the inner AF cells. Therefore, to simplify experimental design and maintain consistency, these NP markers were exclusively assessed in the qPCR analyses in this study.

### 2.7 Protein expression assessment

#### MiRNA transfection

Human IVD cells were transfected with let-7b-5p, miR-100-5p, or a combination of let-7b-5p and miR-100-5p (5 nM of each miRNA in serum-free complete media). The culture media were spiked with the same amount of miRNA mimics after 24 hours. (**Figure 1B(b)**) Conditioned media were collected afterwards, centrifuged at 1,500 rpm for 5 min to remove cellular debris, and stored at -80°C for further analysis.

#### Inflammatory mediator analysis

Fifteen inflammatory mediators of interest were analyzed using the custom Human ProcartaPlex Mix&Match kit (#PPX-15-MXCE7EV) (ThermoFisher Scientific, Toronto, ON, Canada) and measured with the Luminex xMAP system (Serial #MAGPX11188004) (ThermoFisher Scientific, Toronto, ON, Canada) according to the manufacturer’s instructions as previously described.^38^ The samples were pre-diluted 1:3 with DMEM media. Standard curves for each analyte were generated with a 4-parameter logistic (4-PL) algorithm. Both median fluorescence intensity (MFI) corrected with background and the final concentration (pg/mL) were reported by the ThermoFisher ProcartaPlex analysis platform. The concentrations of all analytes were within the detection range with all biological replicates in at least one sample group. Few concentrations were out of the detection range and were extrapolated using the associated standard curves for further analysis.

### 2.8 Statistical analysis and visualization

Statistical analyses were performed by GraphPad Prism 10 (version 10.3.1 (464)) (GraphPad Software, Inc., La Jolla, CA, USA). Shapiro-Wilk normality tests were conducted for all quantitative data.^39^ Two-tailed paired Student’s *t*-test, Welch’s analysis of variance (ANOVA) test, and one-way ANOVA test were used for the parametric data accordingly. Friedman test was used for the nonparametric data. *p* < 0.05 was considered statistically significant. Data are presented as mean ± standard error of the mean (SEM). All assessments were conducted with three or more independent experiments, indicated by “*n*” in the figure legends. Venn diagrams were generated by Venny 2.1 (https://bioinfogp.cnb.csic.es/tools/venny/).

## 3. Results

### 3.1 The miRNA cargo profile and distribution in EVs from human IVD cells

Small RNA sequencing analysis was performed to establish the landscape and evaluate miRNA abundance in IVD cell-derived EVs from tissues of different degrees of degeneration. Three groups were created of non-degenerate (Non-deg), mildly-degenerate (Mildly-deg), and degenerate (Deg) tissues, respectively. A total of 2,632 miRNAs (including isoforms) were identified across the samples and groups, and 14.9% (393 miRNAs) of them were detected with reliable abundance values (count per million, CPM). Of the 393 miRNAs, 184 (46.8%) were shared between the groups (**Figure 2**). Forty-eight miRNAs (12.2%) were detected only in the Non-deg and Mildly-deg groups, 26 miRNAs (6.6%) were detected only in the Non-deg and Deg groups, and 9 miRNAs (2.3%) were detected only in the Mildly-deg and Deg groups (**Figure 2**). In addition, 68 miRNAs (17.3%) were exclusively detected in the Non-deg group, 38 miRNAs (9.7%) were detected solely in the Mildly-deg group, and 20 miRNAs (5.1%) were only detected in the Deg group (**Figure 2**). In summary, less than 50% of miRNAs were shared by all three groups, with the highest number of unique miRNAs found in the Non-deg samples.

**Figure 2.**
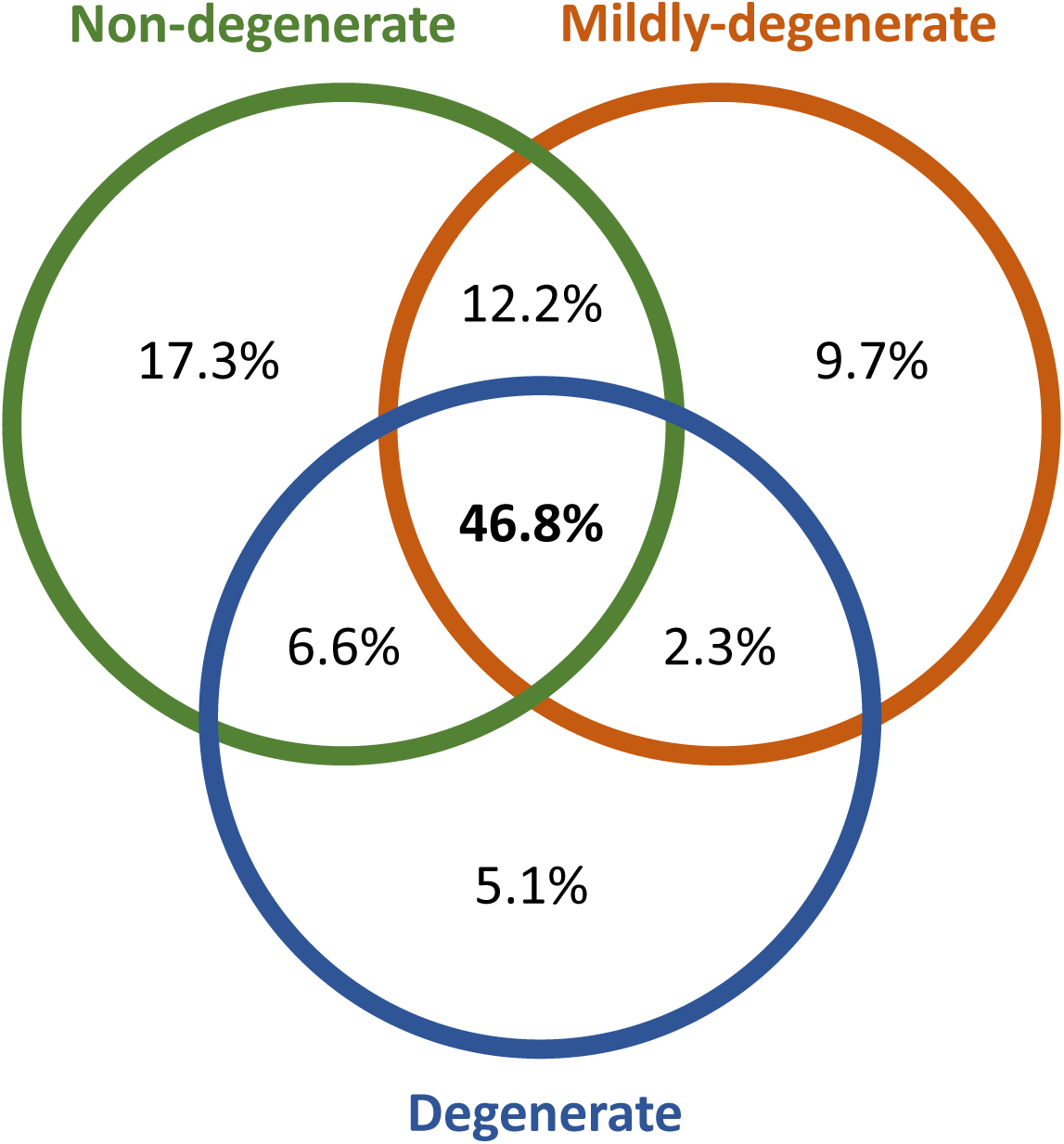
miRNA distribution in EVs isolated from human intervertebral disc (IVD) cells. The Venn diagram represents overlapping and unique miRNAs in EVs from Non-deg (green), Mildly-deg (orange), and Deg (blue) human IVD samples.

### 3.2 Comparisons of the pathways and functional annotations of the miRNAs exclusively detected in one group

The relative expression level of miRNAs exclusively detected in one group was low, ranging between 54 - 714 CPM (**Figure 3A**). Overall, KEGG analysis identified 5 common pathways in the three groups (**Figure 3B** and **Table 3**). The enrichment of pathways such as regulation of the actin cytoskeleton, Hippo signalling, focal adhesion, transforming growth factor (TGF)-beta, and Wnt signalling indicates that the miRNAs are involved in key processes governing cell structure, survival, inflammation, and tissue regeneration. In addition, 5 pathways were exclusively detected in the Deg group and the miRNA cargo of EVs from the Deg samples was associated with the most distinctive KEGG pathways (**Table 3**). The enrichment of pathways such as cell cycle arrest, p53 signalling, vascular endothelial growth factor (VEGF) signalling, apoptosis, and cellular senescence indicates a cellular landscape marked by stress, aging, and degeneration. These pathways are commonly activated in response to DNA damage, oxidative stress, and inflammatory stimuli, all of which are characteristic of IVD degeneration. Their presence suggests that the miRNA cargo within EVs may either reflect a degenerative state or actively modulate these processes, offering potential therapeutic avenues to counteract cell loss, inflammation, and functional decline. A Venn diagram of unique and overlapping GO terms with biological function (BP), cellular component (CC), and molecular function (MF) are in **Figure S1** and **Table S1-3**.

**Figure 3.**
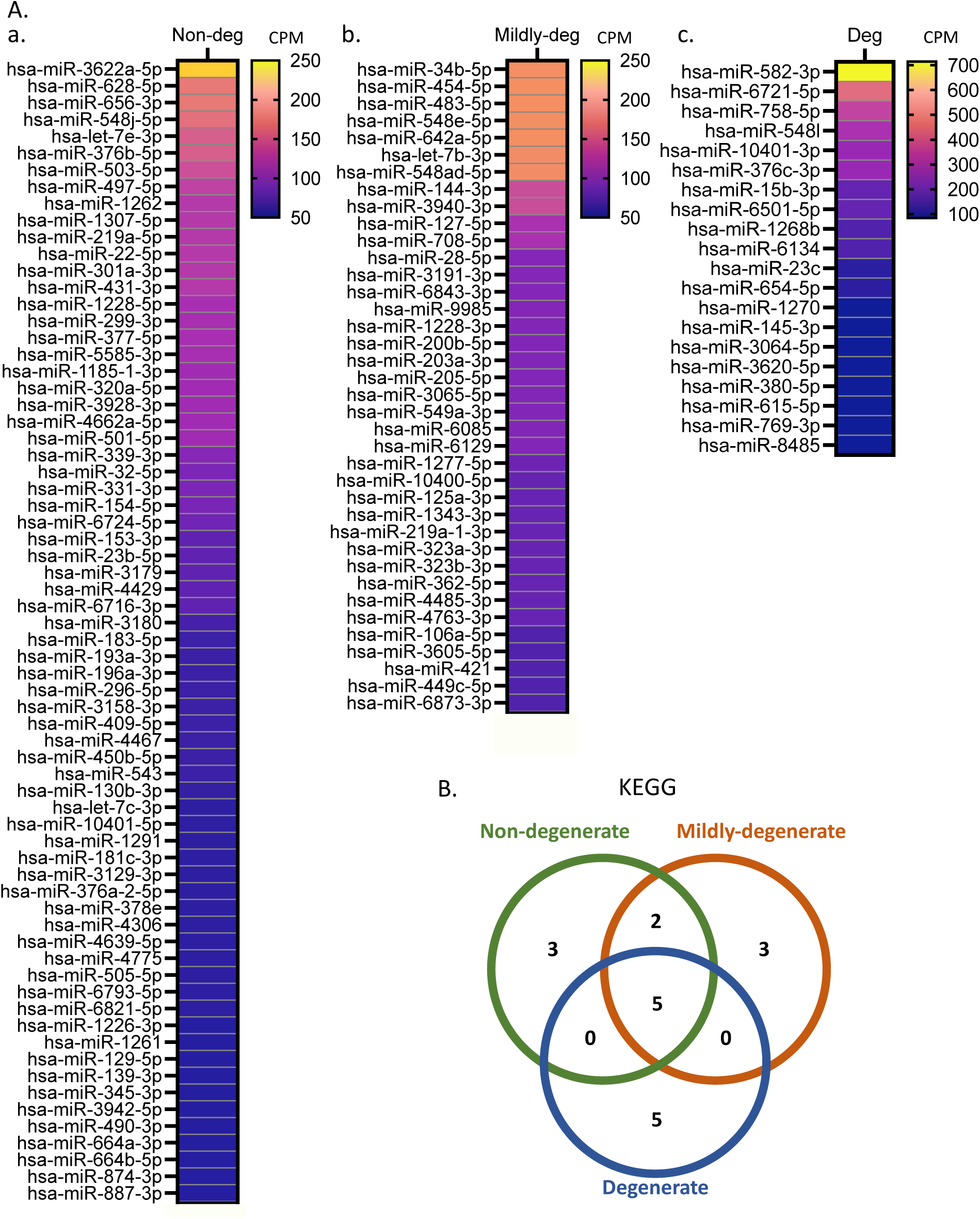
miRNAs and KEGG pathways associated with the uniquely identified miRNAs. **A**. Heatmap demonstrating the expression of miRNAs uniquely detected in the (**a**) Non-deg, (**b**) Mildly-deg, and (**c**) Deg samples. The yellow-blue colour strip represents CPM mean: expression is depicted from high in yellow to low in blue, and rows represent DEMs. *n* = 3-4. **B.** Venn diagram presenting the distribution of the top 10 KEGG pathways associated with musculoskeletal disease, shared and exclusively detected in Non-deg (green), Mildly-deg (orange), and Deg (blue) samples.

**Table 3.** Shared and unique top 10 musculoskeletal disease-associated KEGG pathways of miRNAs detected in the Non-deg, Mildly-deg, and Deg samples.

| Groups | Quantity of shared pathways | Pathways |
| --- | --- | --- |
| Detected in the Non-deg, Mildly-deg, and Deg groups | 5 | Regulation of actin cytoskeleton |
|  |  | Hippo signalling pathway |
|  |  | Focal adhesion |
|  |  | TGF-beta signalling pathway |
|  |  | Wnt signalling pathway |
| Detected in the Non-deg and Mildly-deg groups | 2 | ECM-receptor interaction |
|  |  | PI3K-Akt signalling pathway |
| Exclusively detected in the Non-deg group | 3 | mTOR signalling pathway |
|  |  | AMPK signalling pathway |
|  |  | TNF signalling pathway |
| Exclusively detected in the Mildly-deg group | 3 | FoxO signalling pathway |
|  |  | Adherens junction |
|  |  | MAPK signalling pathway |
| Exclusively detected in the Deg group | 5 | Cell cycle |
|  |  | p53 signalling pathway |
|  |  | VEGF signalling pathway |
|  |  | Apoptosis |
|  |  | Cellular senescence |

### 3.3 Comparisons of the pathways and functional annotations of the shared miRNAs detected in two of the three groups

As with the uniquely detected miRNAs, the shared miRNAs present in two of the three groups were expressed at low levels (**Figure 4**) across all pairwise comparisons: Non-deg and Mildly-deg groups (**Figure 4A**; 54-546 CPM), Non-deg and Deg groups (**Figure 4B**; 54-571 CPM), and Mildly-deg and Deg groups (**Figure 4C**; 78-367 CPM). KEGG analysis consistently highlighted focal adhesion and phosphatidylinositol 3-kinase (PI3K)-Akt signalling pathway in the comparisons (**Table 4**). The pathways integrate cell-ECM mechanotransduction with survival and anabolic control, supporting proteoglycan and collagen synthesis while constraining inflammatory cascades and apoptosis. Their recurrence suggests common, stage-spanning activities through which EV-miRNAs modulate disc homeostasis. The unique pathway and annotation results are described in the *Supplementary Results* (**Table S4-12**). Briefly, the pairwise analyses reveal that shared pathways focal adhesion and PI3K-Akt signalling pathway span stages, early Non-deg-anchored shifts in growth control and matrix signalling, Mildly-deg-anchored stress and phenotype reprogramming, and Deg-anchored cytoskeletal and mechanical rewiring with metabolic and junctional liabilities. This framework pinpoints actionable pathways for EV-miRNA interventions aimed at restoring mechanotransduction balance, sustaining anabolic capacity, and curbing inflammation, apoptosis, and neovascular cues.

**Figure 4.**
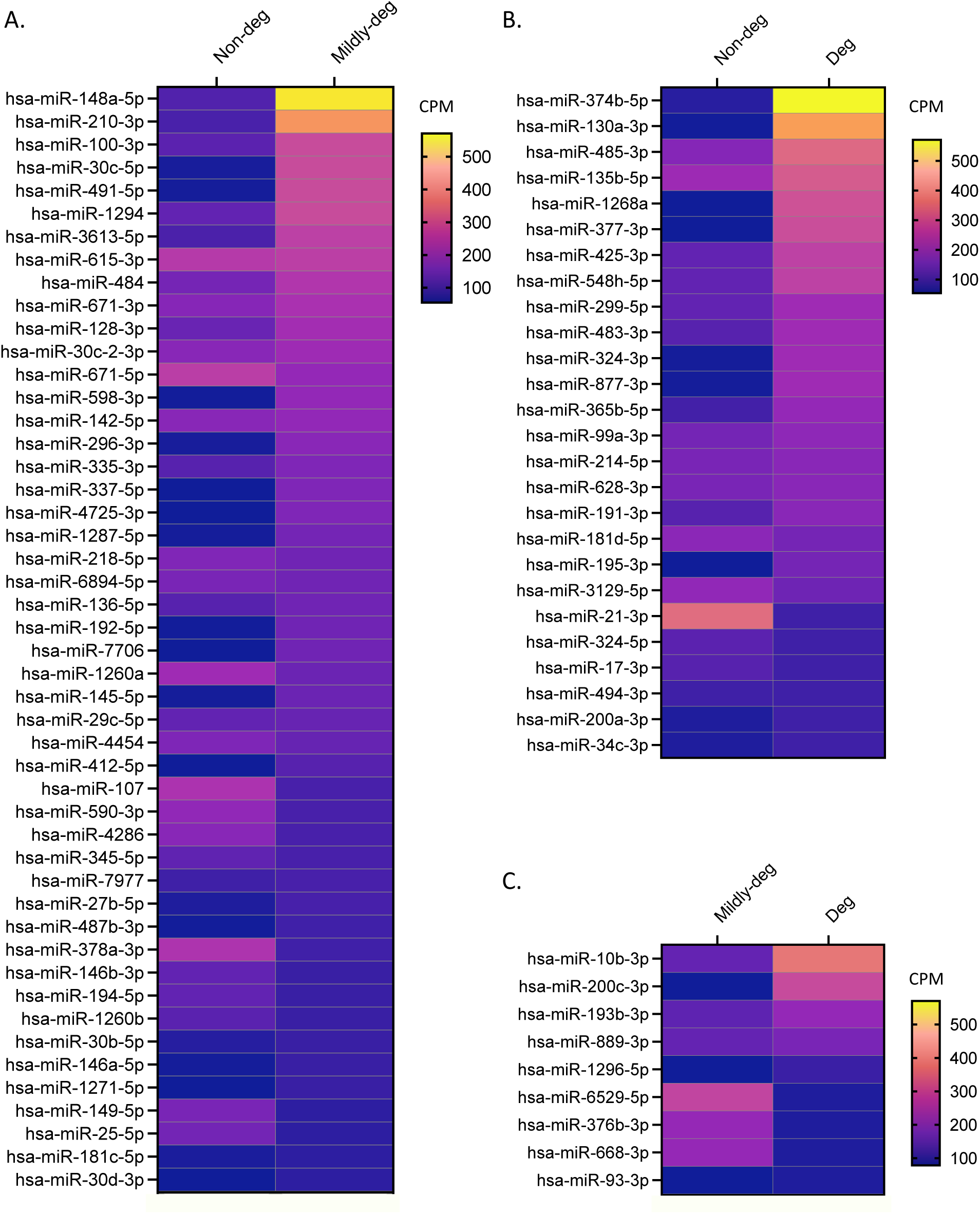
The relative expression of miRNAs shared between two of the three groups. The Heatmap shows differentially expressed miRNAs comparing (**A**) Non-deg and Mildly-deg, (**B**) Non-deg and Deg, and (**C**) Mildly-deg and Deg samples. The yellow-blue colour strip represents CPM mean: expression is depicted from high in yellow to low in blue, and rows represent DEMs. *n* = 3-4.

**Table 4.**
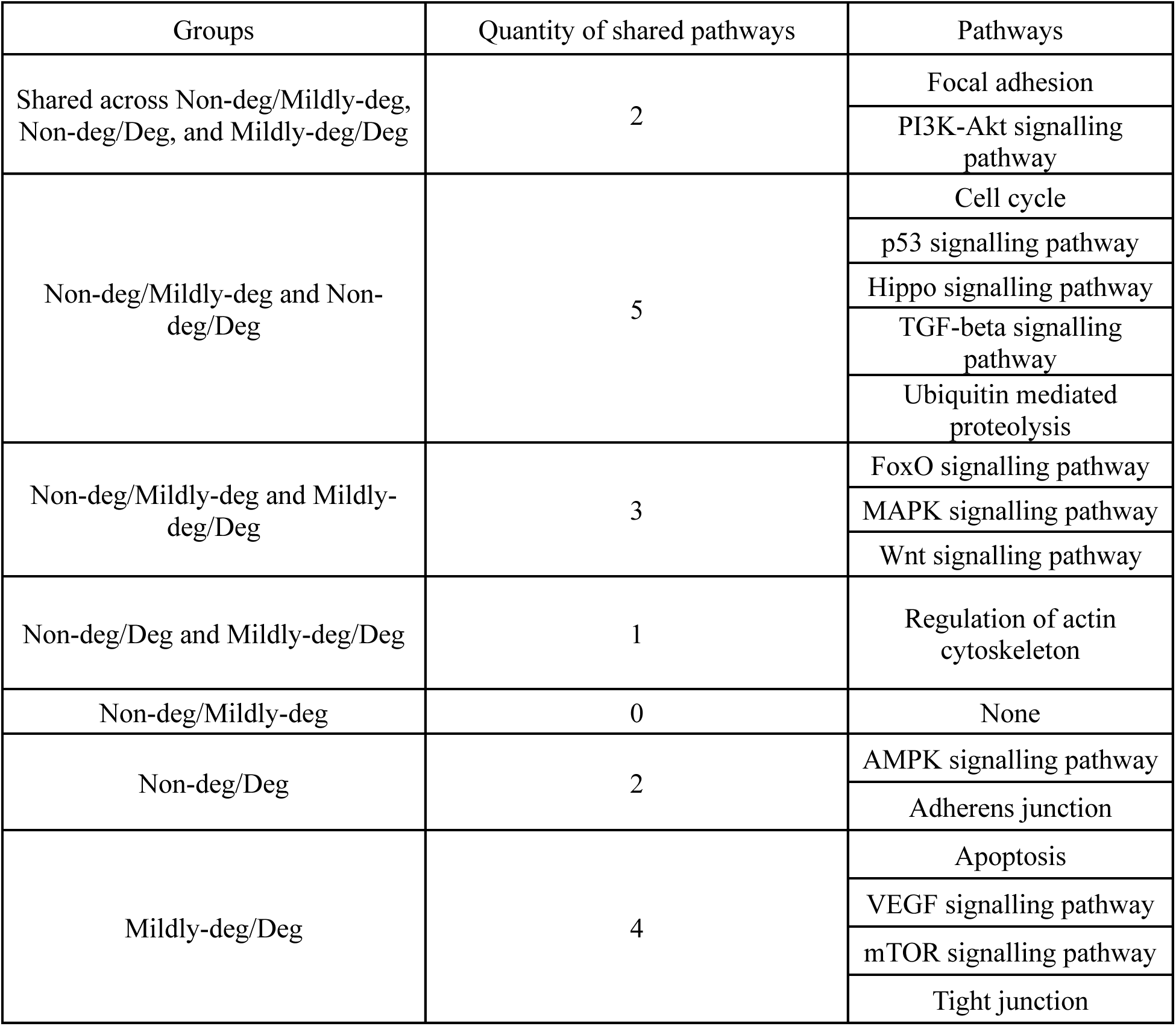
Shared and unique top 10 musculoskeletal disease-associated KEGG pathways of miRNAs detected in two of the three groups.

### 3.4 The differential expression and comparisons of the pathways and functional annotations of the miRNAs shared between all groups

Across the 184 miRNAs shared by all groups, the expression level ranged from 54-227,602 CPM. KEGG analysis converged on pathways governing senescence control, survival/anabolism, and mechanotransduction (**Figure 5A**). Consistent with this, GO-BP highlighted cell cycle/protein phosphorylation and ECM organization (**Figure S2A**); GO-CC emphasized focal adhesions, cytoskeleton, and ECM/collagen compartments (**Figure S2B**); and GO-MF underscored integrin/cadherin/collagen binding alongside kinase and ubiquitin-transferase activities (**Figure S2C**).

**Figure 5.**
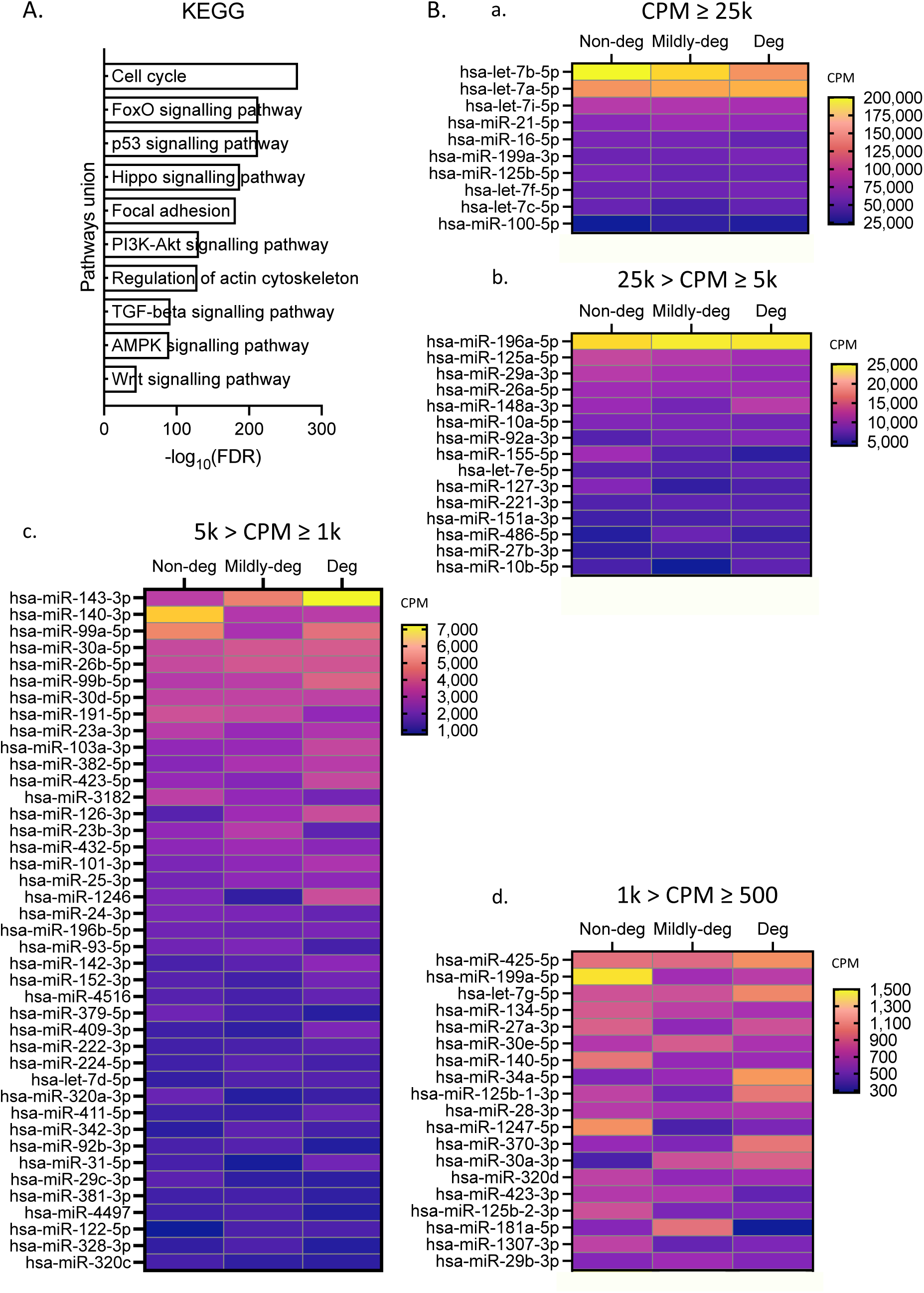
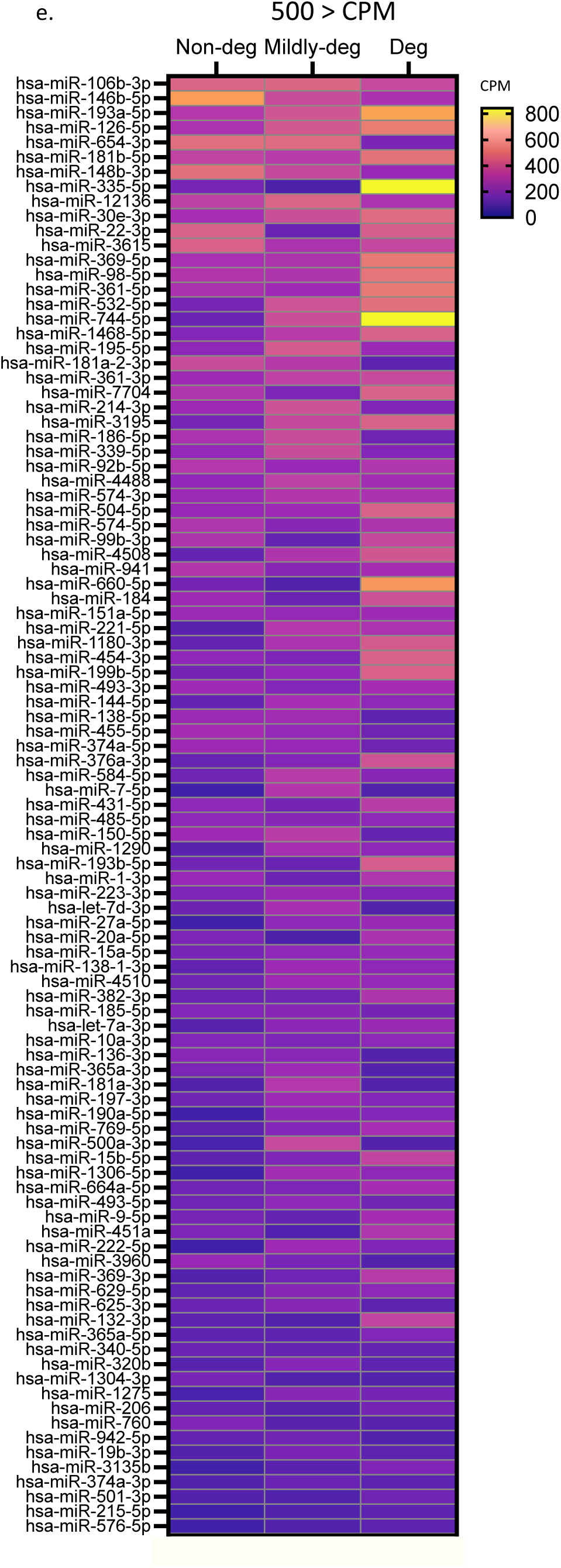
KEGG analysis and relative expression of miRNA shared between all three groups. (**A**) KEGG analyses showing the top 10 musculoskeletal disease-associated pathways, *p*-value threshold of 0.05. The x-axis represents the logarithmic scale of the FDR (-Log10 (FDR)), and the y-axis describes the enriched KEGG pathways. (**B**) Heatmaps displaying miRNAs within different CPM ranges: **(a)**, CPM ≥ 25k, **(b)**, 25k > CPM ≥ 5k, **(c)**, 5k > CPM ≥ 1k, **(d)**, 1k > CPM ≥ 500, and **(e)**, 500 > CPM. The yellow-blue colour strip represents CPM mean: expression is depicted from high in yellow to low in blue, and rows represent DEMs. *n* = 3-4.

Ten miRNAs were expressed at very high levels exceeding 25,000 CPM. The top 10 abundant miRNAs are let-7b-5p, let-7a-5p, let-7i-5p, miR-21-5p, miR-16-5p, miR-199a-3p, miR-125b-5p, let-7f-5p, let-7c-5p, and miR-100-5p (**Figure 5B**). KEGG analysis supports the shared core: cell cycle, forkhead box O (FoxO), Hippo, and p53 signalling pathways, indicating a strong control of senescence checkpoints, survival, and mechanotransduction. The added terms: proteoglycans in cancer, adherens junction, and oocyte meiosis expand their predicted functions toward broad stress-response and adhesion remodelling. GO-BP/CC/MF reinforce a transcription-centric program anchored in the nucleus/nucleoplasm and coupled to cytoskeletal compartments. (**Figure S3A**)

Fifteen miRNAs were expressed at high levels with a CPM range of 5,000 - 25,000 (**Figure 5B**). KEGG analysis largely supported the shared backbone: focal adhesion, PI3K-Akt, Hippo, actin cytoskeleton, TGF-beta, Wnt, FoxO signalling pathways and uniquely adds mitogen-activated protein kinase (MAPK), adherens junction, and ubiquitin-mediated proteolysis, linking survival/anabolism to junctional rewiring and proteostasis. GO terms concentrate on ECM organization and focal/adherens junction locales with integrin/cadherin/collagen binding and kinase/ubiquitin activities. (**Figure S3B**) Forty-one miRNAs were expressed at moderate levels with a CPM range of 1,000 - 5,000 (**Figure 5B**). KEGG retains core pathways and uniquely introduces ECM-receptor interaction, MAPK, and ubiquitin-mediated proteolysis, emphasizing matrix-receptor crosstalk and stress adaptation. GO highlights basement membrane/ECM compartments and SMAD/actin/collagen binding. (**Figure S3C**) Nineteen miRNAs were expressed at low levels with a CPM mean range of 500 - 1,000 (**Figure 5B**). KEGG overlaps with the core and uniquely adds ECM-receptor interaction, adherens junction, MAPK, and VEGF signalling pathways, pointing to adhesion remodelling and angiogenic cues. GO emphasizes cytoskeleton/focal adhesion/ECM locales with SMAD/actin and ubiquitin/kinase interfaces. (**Figure S3D**) Ninety-nine miRNAs were expressed at very low levels with a CPM range of 54 - 500 (**Figure 5B**). KEGG again aligns with the shared backbone and adds ECM-receptor interaction, adherens junction, and MAPK signalling pathways, consistent with low-abundance modulators of adhesion-cytoskeleton-inflammation coupling. GO terms support the functions (**Figure S3E**).

All abundance groups converge on cellular-senescence control and survival/anabolism and on mechanotransduction/adhesion-ECM circuitry. Together, the data support a model in which EV-miRNAs cooperatively sustain matrix homeostasis, limit apoptosis and senescence, and dampen inflammatory and angiogenic drivers across degeneration stages.

### 3.5 Selection of five miRNAs for functional testing

We focused on miRNAs with high abundance across samples, using the average abundance of all groups as a criterion for selection (**Table 5**). Let-7b-5p and let-7a-5p were the two most abundant miRNAs identified in our analysis and are known for anti-inflammatory effects in various models.^40-44^ Notably, let-7b-5p exhibited a significantly lower expression in Non-deg samples than in Deg samples (*p* = 0.0243). Let-7f-5p showed consistent expression across all groups, as indicated by its small standard deviation of CPM values. MiR-16-5p supplementation has shown mixed effects on inflammatory responses in osteoarthritis and spinal cord injury studies^45-47^, and it showed a decreasing trend in expression from Non-deg to Deg samples (*p* = 0.0390). In addition, miR-100-5p was evenly expressed across the groups, and was selected because it is known primarily as a tumor suppressor in various cancers and could potentially play a role in the regulation of cellular senescence.^48,49^ In summary, our selected miRNAs include 1) three with a similar expression level and a potential anti-inflammatory effect (let-7b-5p, let-7a-5p, and let-7f-5p), 2) one with differential expression potentially linked to inflammatory responses (miR-16-5p), and 3) one implicated in the regulation of cellular senescence (miR-100-5p) (**Figure 6A and Table 5**).

**Figure 6.**
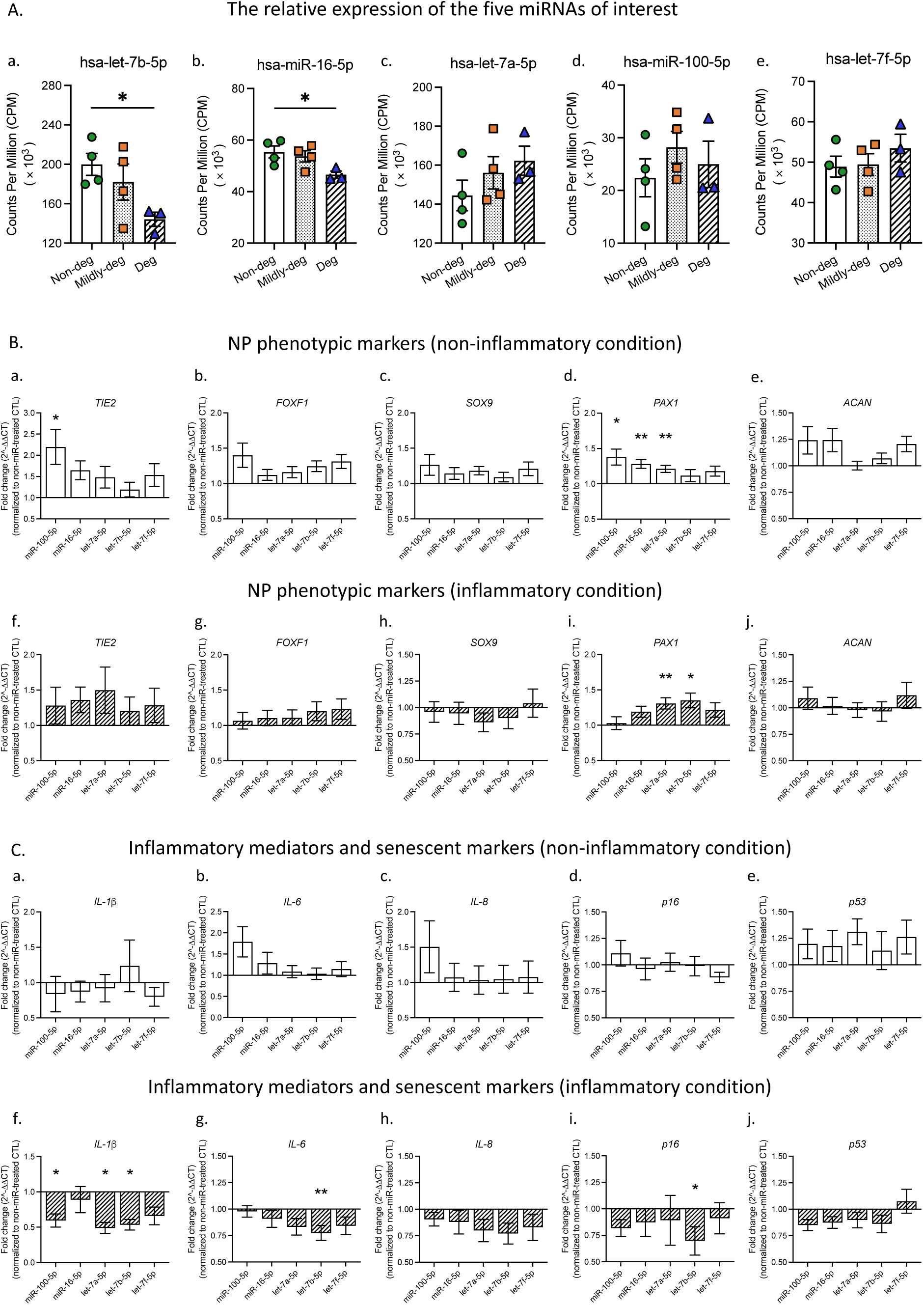
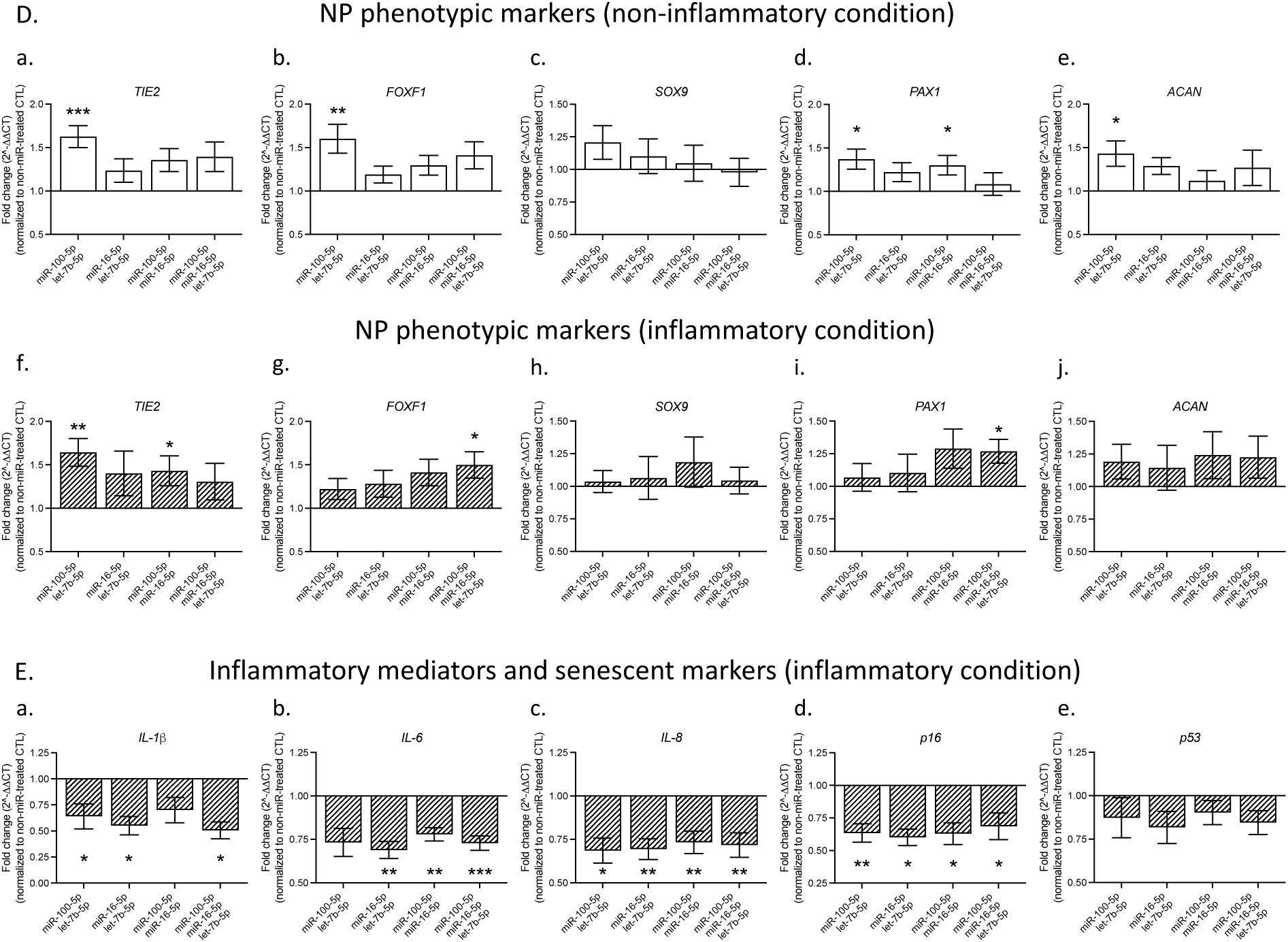
The relative expression and regulatory effects of five selected miRNAs. **(A)** CPM. Regulatory effects of single miRNA mimic treatment on NP cell phenotypic marker expression: under (**B**) non-inflammatory conditions and (**C**) inflammatory conditions. Regulatory effects of single miRNA mimic treatment on inflammatory mediators and senescence marker expression: under (**D**) non-inflammatory conditions and (**E**) inflammatory conditions. Regulatory effects of miRNA mimic combination treatment on (**F)** NP cell phenotypic markers and (**G)** inflammatory mediators and senescence markers. Data were normalized to vehicle-treated cells. All values are presented as mean ± SEM. The data in **A** was analyzed by Welch’s ANOVA: *p* < 0.05. *n* = 3-4. The data in **B** *(SOX9*, *IL-6*, and *IL-8)* and **C** (*p16*) are non-parametric and were analyzed by Friedman test with Dunn’s multiple comparisons test as indicated in *Methods*. *, **, and *** indicate a statistical significance of *p* < 0.05, *p* < 0.01, and *p* < 0.001. All other data were analyzed by one-way ANOVA with Dunnett’s T3 multiple comparisons test. *n* = 10.

**Table 5.** Top 10 abundant shared miRNAs in the Non-deg, Mildly-deg, and Deg samples.

| Abundance rank | Overall | Non-deg group | Mildly-deg group | Deg group |
| --- | --- | --- | --- | --- |
| 1 | **let-7b-5p** | let-7b-5p | let-7b-5p | let-7a-5p |
| 2 | **let-7a-5p** | let-7a-5p | let-7a-5p | let-7b-5p |
| 3 | let-7i-5p | let-7i-5p | let-7i-5p | let-7i-5p |
| 4 | miR-21-5p | miR-21-5p | miR-21-5p | miR-21-5p |
| 5 | **miR-16-5p** | miR-16-5p | miR-16-5p | miR-199a-3p |
| 6 | miR-199a-3p | miR-125b-5p | miR-199a-3p | let-7f-5p |
| 7 | miR-125b-5p | miR-199a-3p | let-7f-5p | miR-125b-5p |
| 8 | **let-7f-5p** | let-7f-5p | miR-125b-5p | miR-16-5p |
| 9 | let-7c-5p | let-7c-5p | let-7c-5p | let-7c-5p |
| 10 | **miR-100-5p** | miR-196a-5p | miR-100-5p | miR-100-5p |

Bold font indicates miRNAs of interest.

### 3.6 The regulatory effect of the selected miRNAs in human IVD cells

#### 3.6.1 The regulatory effects of single-miRNA treatment on cell phenotype

Each of the five selected miRNA mimics was applied to IVD cells under non-inflammatory and inflammatory conditions (**Figure S4A**) and the expression of NP markers was assessed. MiR-100-5p significantly upregulated *TIE2* expression (2.20 ± 0.42, *p* = 0.0201) under non-inflammatory conditions (**Figure 6B**). *Paired Box 1* (*PAX1)* expression was significantly upregulated by miR-100-5p (1.38 ± 0.11, *p* = 0.0125), miR-16-5p (1.28 ± 0.06, *p* = 0.0035), and let-7a-5p (1.21 ± 0.05, *p* = 0.0079). No significant change was observed in the expression of the NP markers *Forkhead Box F1 (FOXF1)*, *SRY-Box Transcription Factor 9* (*SOX9)*, and *ACAN* under non-inflammatory conditions. *PAX1* expression was significantly upregulated by let-7a-5p (1.31 ± 0.08, *p* = 0.0074) and let-7b-5p (1.35 ± 0.11, *p* = 0.0308) under inflammatory conditions. In addition, no significant change was observed in the expression of NP markers *TIE2*, *FOXF1*, *SOX9*, and *ACAN* under inflammatory conditions (**Figure 6B**).

#### 3.6.2 The regulatory effect of single-miRNA treatment on inflammatory mediators and senescence markers

Inflammatory mediator (*Interleukin (IL)-1 beta (IL-1ß)*, *IL-6*, and *IL-8*) and senescence marker (*p16* and *p53*) expression was very low under non-inflammatory conditions, and no significant change was observed in their expression following miRNA treatment (**Figure 6C**). Their expression was as expected elevated under inflammatory conditions (**Figure S4A**), where miR-100-5p (0.59 ± 0.09, *p* = 0.0382), let-7a-5p (0.49 ± 0.08, *p* = 0.0107), and let-7b-5p (0.53 ± 0.08, *p* = 0.0144) significantly downregulated *IL-1ß*. Let-7b-5p significantly downregulated *IL-6* (0.77 ± 0.07, *p* = 0.0094) and *p16* (0.70 ± 0.13, *p* = 0.0461). No significant change was observed in the expression of *IL-8* and *p53* (**Figure 6C**).

We selected miR-100-5p, let-7b-5p, and miR-16-5p based on their regulatory effects on key NP markers, inflammatory mediators, and senescence markers under inflammatory and non-inflammatory conditions.

#### 3.6.3 The regulatory effects of miRNA combination treatment on cell phenotype

To explore their combined effects, miRNA mimics were combined in four treatment groups: 1) miR-100-5p + let-7b-5p, 2) miR-16-5p + let-7b-5p, 3) miR-100-5p + miR-16-5p, and 4) miR-100-5p + miR-16-5p + let-7b-5p. These treatments were applied to IVD cells under inflammatory and non-inflammatory conditions, and their effects on NP markers, inflammatory mediators, and senescence markers were assessed. MiR-100-5p + let-7b-5p significantly upregulated the expression of *TIE2* (1.63 ± 0.13, *p* = 0.0005), *FOXF1* (1.60 ± 0.17, *p* = 0.0060), *PAX1* (1.37 ± 0.12, *p* = 0.0211) and *ACAN* (1.43 ± 0.15, *p* = 0.0308) under non-inflammatory conditions. No significant change was observed in the expression of the NP marker *SOX9* under non-inflammatory conditions. MiR-100-5p + let-7b-5p significantly upregulated *TIE2* expression (1.64 ± 0.16, *p* = 0.0089) under inflammatory conditions. MiR-100-5p + miR-16-5p + let-7b-5p significantly upregulated *FOXF1* (1.50 ± 0.15, *p* = 0.0173) and *PAX1* (1.27 ± 0.09, *p* = 0.0350) expression under inflammatory conditions. No significant changes were observed in the expression of the NP markers *SOX9* and *ACAN* under inflammatory conditions (**Figure 6D)**.

#### 3.6.4 The regulatory effects of miRNA combination treatment on inflammatory mediators and senescence markers

Under inflammatory conditions, miR-100-5p + let-7b-5p significantly downregulated the expression of *IL-1ß* (0.64 ± 0.12, *p* = 0.0338), *IL-8* (0.69 ± 0.07, *p* = 0.0487), and *p16* (0.63 ± 0.07, *p* = 0.0079). MiR-16-5p + let-7b-5p significantly downregulated the expression of *IL-1ß* (0.55 ± 0.09, *p* = 0.0107), *IL-6* (0.69 ± 0.05, *p* = 0.0024), *IL-8* (0.69 ± 0.06, *p* = 0.0024), and *p16* (0.60 ± 0.06, *p* = 0.0167). MiR-100-5p + miR-16-5p significantly downregulated the expression of *IL-6* (0.78 ± 0.04, *p* = 0.0019), *IL-8* (0.73 ± 0.07, *p* = 0.0045), and *p16* (0.63 ± 0.08, *p* = 0.0113). In addition, miR-100-5p + miR-16-5p + let-7b-5p significantly downregulated the expression of *IL-1ß* (0.51 ± 0.08, *p* = 0.0200), *IL-6* (0.73 ± 0.04, *p* = 0.0007), *IL-8* (0.72 ± 0.07, *p* = 0.0058), and *p16* (0.69 ± 0.10, *p* = 0.0394) (**Figure 6E**). No significant changes were observed in the expression of *p53* (**Figure 6E**) under inflammatory conditions and all the markers under non-inflammatory conditions (**Figure S4B**).

Overall, the results suggest that the combination treatment with miR-100-5p + let-7b-5p have promising regulatory effects on maintaining IVD cell phenotype with anti-inflammatory and anti-senescence effects in inflammatory conditions.

### 3.7 The anti-inflammatory effects of miRNA treatment at the protein level

We quantified the expression of 15 inflammatory mediators under inflammatory conditions (**Figure S5**). Our results showed that the production of three cytokines (IL-6, IL-1 alpha (IL-1α), and tumor necrosis factor alpha (TNF-α)) was significantly reduced by miR-100-5p, let-7b-5p, and their combination treatment. In addition, the production of two cytokines (IL-1ß and interferon gamma (IFN-γ)) was significantly reduced by the combination treatment of miR-100-5p and let-7b-5p. A decreasing trend for IL-21 was observed following all treatments; however, no statistically significant difference was detected. Anti-inflammatory mediators were also assessed. IL-10 expression was significantly increased by miR-100-5p, let-7b-5p, and their combination treatment compared with the control group (**Figure 7A**). An increasing trend of IL-4 production was observed following all treatments; however, no statistically significant difference was detected (**Figure 7A**). Quantitative data are presented in **Table S13**.

**Figure 7.**
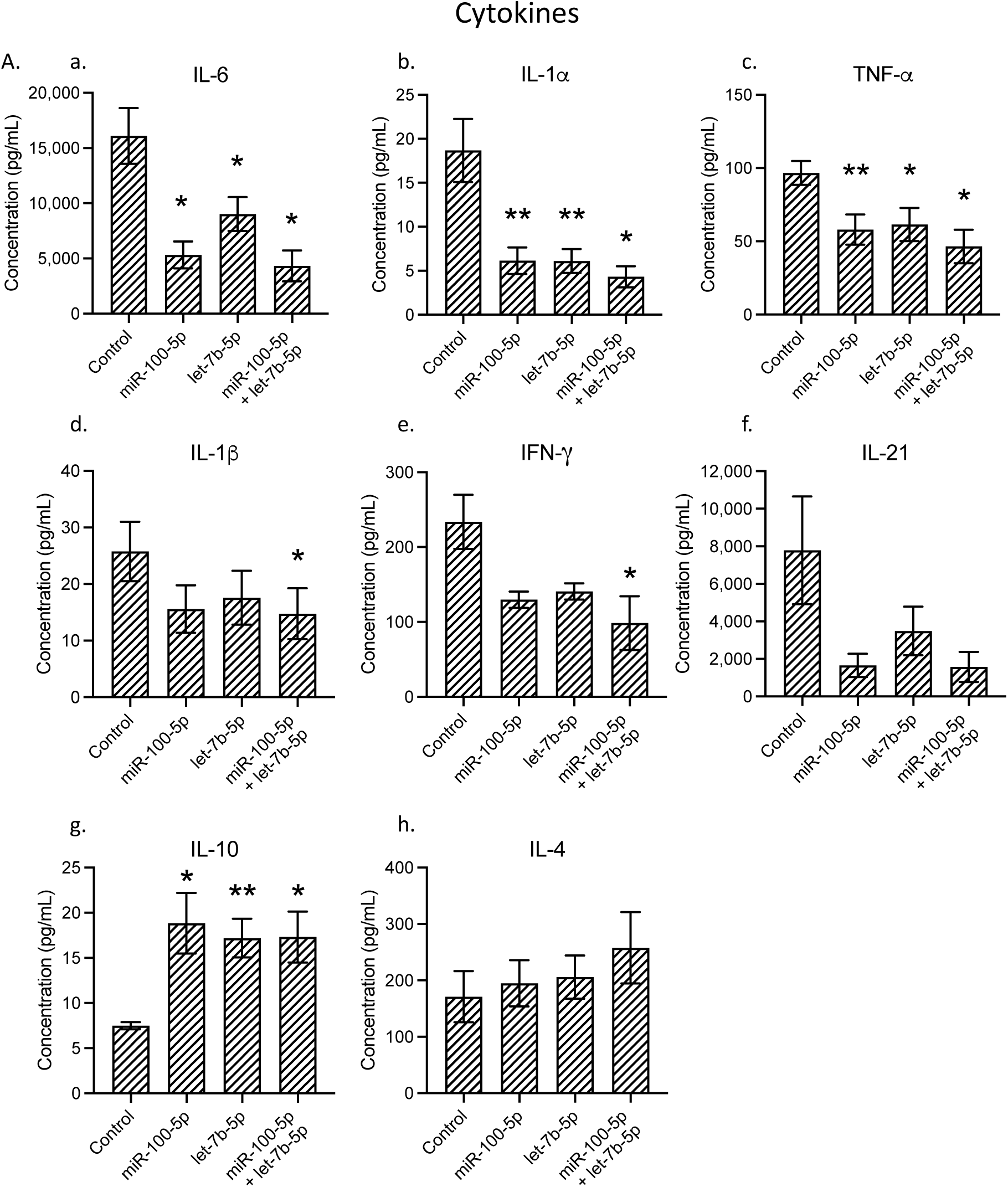
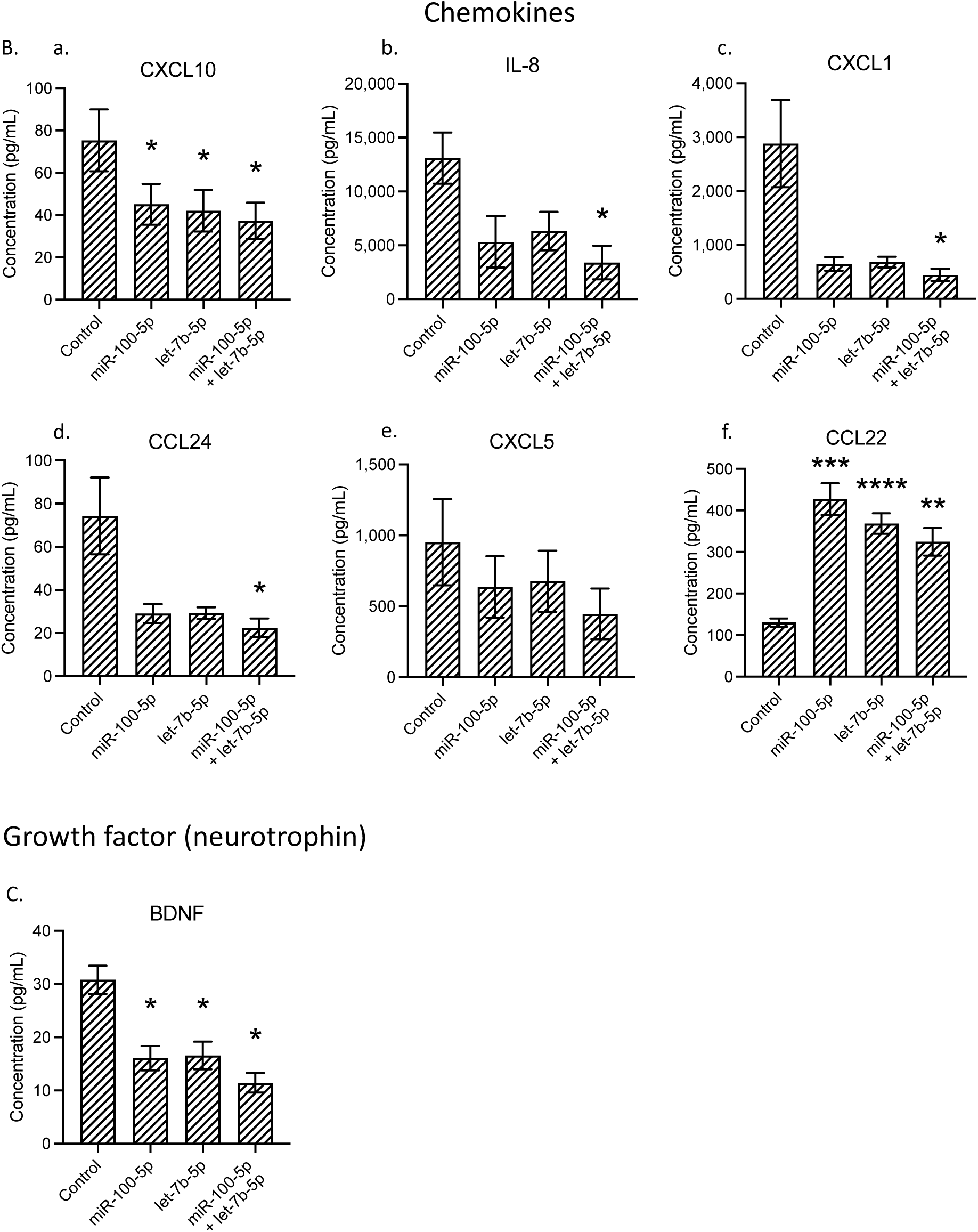
Protein expression of inflammatory mediators post miRNA treatment. (**A**) cytokine expression (**B**) chemokine expression, and (**C**) growth factor expression. Concentrations are presented as mean ± SEM in pg/mL. All data were analyzed by one-way ANOVA with Tukey’s multiple comparisons test. *, **, ***, and **** indicate a statistical significance of *p* < 0.05, *p* < 0.01, *p* < 0.001, and *p* < 0.0001, compared with the control group, respectively. *n* = 8.

In addition, chemokine CXCL10 expression was significantly decreased by miR-100-5p, let-7b-5p, and their combination treatment compared with the control group. Furthermore, the production of three chemokines (IL-8, CXCL1, and CCL24) was significantly reduced by the combination treatment of miR-100-5p and let-7b-5p. A decreasing trend for CXCL5 was observed following all treatments; however, no statistically significant difference was detected. The expression of anti-inflammatory chemokine CCL22 was significantly increased by miR-100-5p, let-7b-5p, and their combination treatment compared with the control group (**Figure 7B**). The expression of growth factor brain-derived neurotrophic factor (BDNF) was significantly decreased by miR-100-5p, let-7b-5p, and their combination treatment compared with the control group (**Figure 7C**). Quantitative data are presented in **Table S13**.

## 4 Discussion

This work combines insights from profiling miRNA cargo in EVs with functional prediction analyses to identify miRNAs with specific roles in IVD maintenance and degeneration. Previous studies emphasized the significance of miRNAs in IVD degeneration^11,12,50,51^ and their emerging roles in EV-mediated cell communication, which can potentially be leveraged for regenerative medicine.^21,52-54^ However, while studies have profiled miRNAs in IVD tissues^55-57^, there is a lack of systematic investigation into miRNA cargo in EVs derived from human IVD cells, especially across different stages of degeneration. This gap motivates our study’s objectives: to profile miRNA cargo in EVs from human IVD cells representing various degeneration grades and to explore the therapeutic functions of selected miRNAs through function prediction-guided screening.

Our results showed that 46.8% of the miRNA cargo was shared among Non-deg, Mildly-deg, and Deg samples. This is much less than the proportion of shared protein cargo from the same EV samples, which was 88.6%^23^, suggesting a more dynamic role for miRNAs in response to degeneration-specific cellular changes. Specific miRNAs, such as miR-100-5p and let-7b-5p, were identified as highly enriched in EVs from IVD cells. These miRNAs demonstrated distinct regulatory functions, including the modulation of NP phenotypic markers, inflammatory mediators, and senescence markers. For instance, miR-100-5p increased *TIE2* expression, which is essential for cell survival, proliferation, cellular maintenance, and regenerative responses in IVD cells.^58-60^ At the same time, let-7b-5p reduced *IL-1ß* expression, a pro-inflammatory cytokine implicated in inflammatory responses, cell apoptosis, cellular senescence, and ECM breakdown within degenerated IVD tissues.^61,62^ Moreover, a combination of these miRNAs demonstrated synergistic therapeutic effects on promoting IVD cell phenotypic marker expression and reducing inflammatory mediator and senescence marker expression, suggesting potential for combined miRNA therapies in future applications.

### 4.1 Comparison of EV-enriched and tissue lysate-enriched miRNAs in human IVD

Previous studies have profiled miRNAs from human IVD tissue (**Table 6**). Ohrt-Nissen *et al.* compared miRNA profiles of Deg NP and AF tissues and highlighted a high abundance of growth factor-associated signalling pathways influenced by miRNAs, including TGF-beta, platelet-derived growth factor, insulin-like growth factor, epidermal growth factor signalling pathways, which are critical for cellular processes in IVD homeostasis.^55^ Zhao *et al.* profiled and compared miRNAs in Non-deg and Deg NP tissue, linking the dysregulated miRNAs to anti-apoptosis, cell proliferation, cell survival, and angiogenesis-associated pathways, suggesting their role in the homeostasis and pathogenesis of IVD degeneration.^56^ In addition, Sherafatian *et al.* conducted a meta-analysis of three miRNA datasets of Non-deg and Deg NP tissue, identifying commonly dysregulated miRNAs associated with anti-apoptosis and cell proliferation-associated pathways.^57^

**Table 6.**
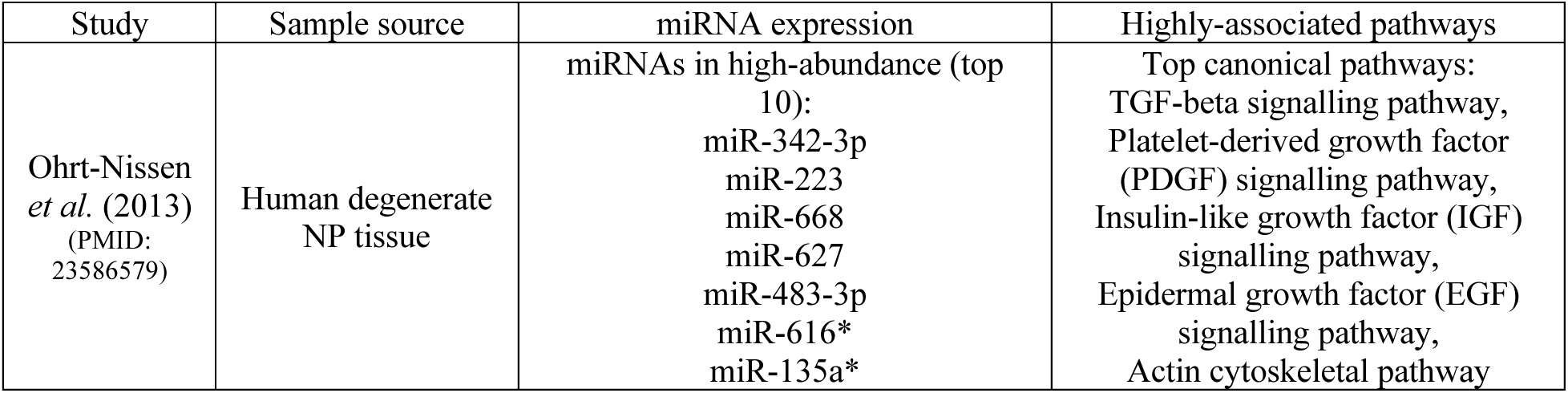
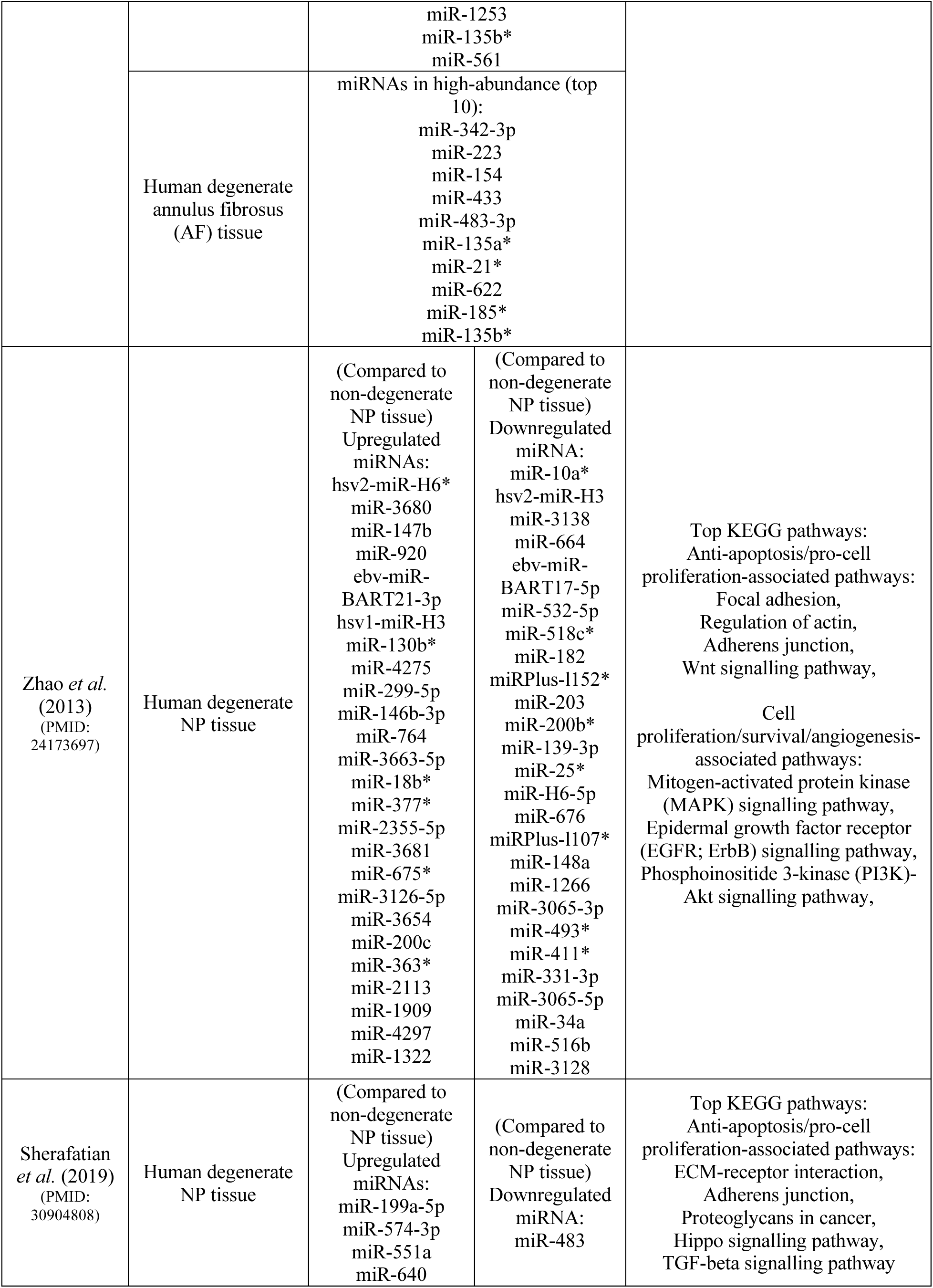
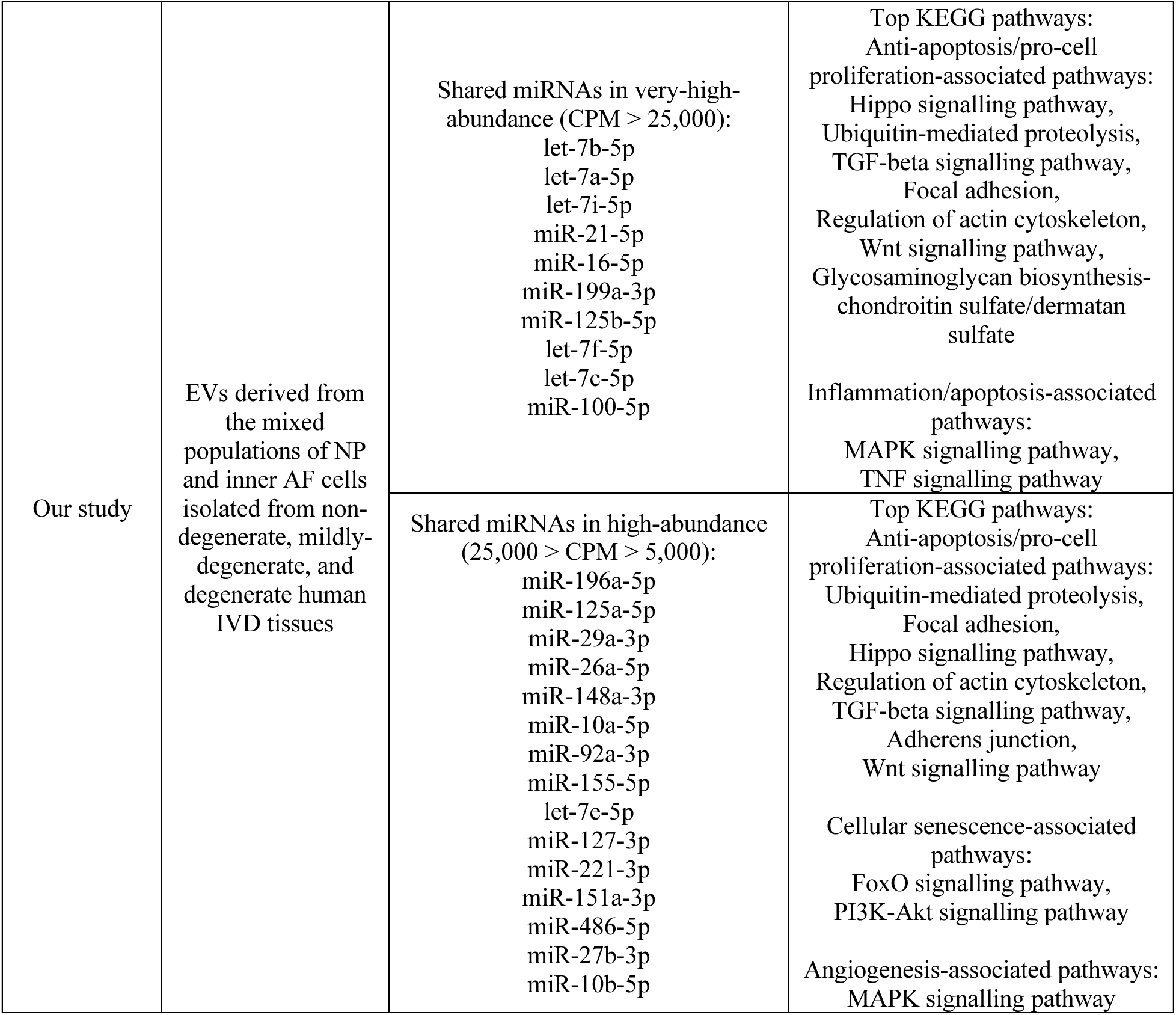
Comparison of EV-enriched and tissue lysate-enriched miRNAs in human IVD.

Our findings reveal that the miRNA profiles of EVs derived from IVD cells differ significantly from those of tissue lysates. We did not identify any overlap between miRNAs with an abundance above 5,000 CPM in our study with the high-abundance or dysregulated miRNA described in the studies mentioned above.^55-57^ However, pathways associated with anti-apoptosis, cell proliferation, angiogenesis, inflammation, and cellular senescence were also identified in our samples. These differences may indicate the selective packaging of miRNAs into EVs, aligning with studies demonstrating that EVs actively transport specific miRNAs^63-65^, likely reflecting the regulatory needs of target cells under specific physiological and pathological conditions. In addition, when interpreting the different miRNA profiles between EV and tissue samples, parameters such as tissue source and technical assessment should be taken into consideration. We purified miRNAs from EVs derived from a mixed population of NP and inner AF cells isolated from human IVD tissues with different degrees of degeneration, while the other studies purified miRNAs directly from Non-deg and Deg human NP and AF tissues and not from EVs, respectively. In addition, we evaluated EV-enriched miRNA profiles using the small RNA sequencing technique, while the other studies used microarray assays. Future studies comparing miRNA profiles between EV and cell/tissue lysate samples should use the same culture system and analytical technique and method to minimize systematic variations.

### 4.2 Comparison of human IVD cell-derived EV-enriched miRNAs and NP stem cell-derived exosome-enriched miRNAs

A previous study reported relative expression of miRNAs from exosomes derived from human NP-derived stem cells (NPSC) isolated from Non-deg and Deg NP tissues (**Table 7**).^66^ We detected all ten significantly dysregulated miRNAs that were reported and identified different relative expression patterns. Particularly, the authors detected higher expression of miR-100-5p compared with that of let-7b-5p. Their relative expression significantly increased in the Deg samples. We detected the highest average expression of let-7b-5p over all detected miRNAs. The average expression of miR-100-5p was ranked tenth among all detected miRNAs in our samples. In addition, we found that the relative expression of let-7b-5p significantly decreased in the Deg samples compared with that of the Non-deg samples, which showed the opposite expression pattern to the reported data. We did not detect a significant difference in the relative expression of miR-100-5p among our samples, which is also different from the reported data.^66^ The remaining eight reported miRNAs were also detected in all samples with various expression levels, except for miR-107, which was detected only in the Non-deg and Mildly-deg samples in our study.

**Table 7.**
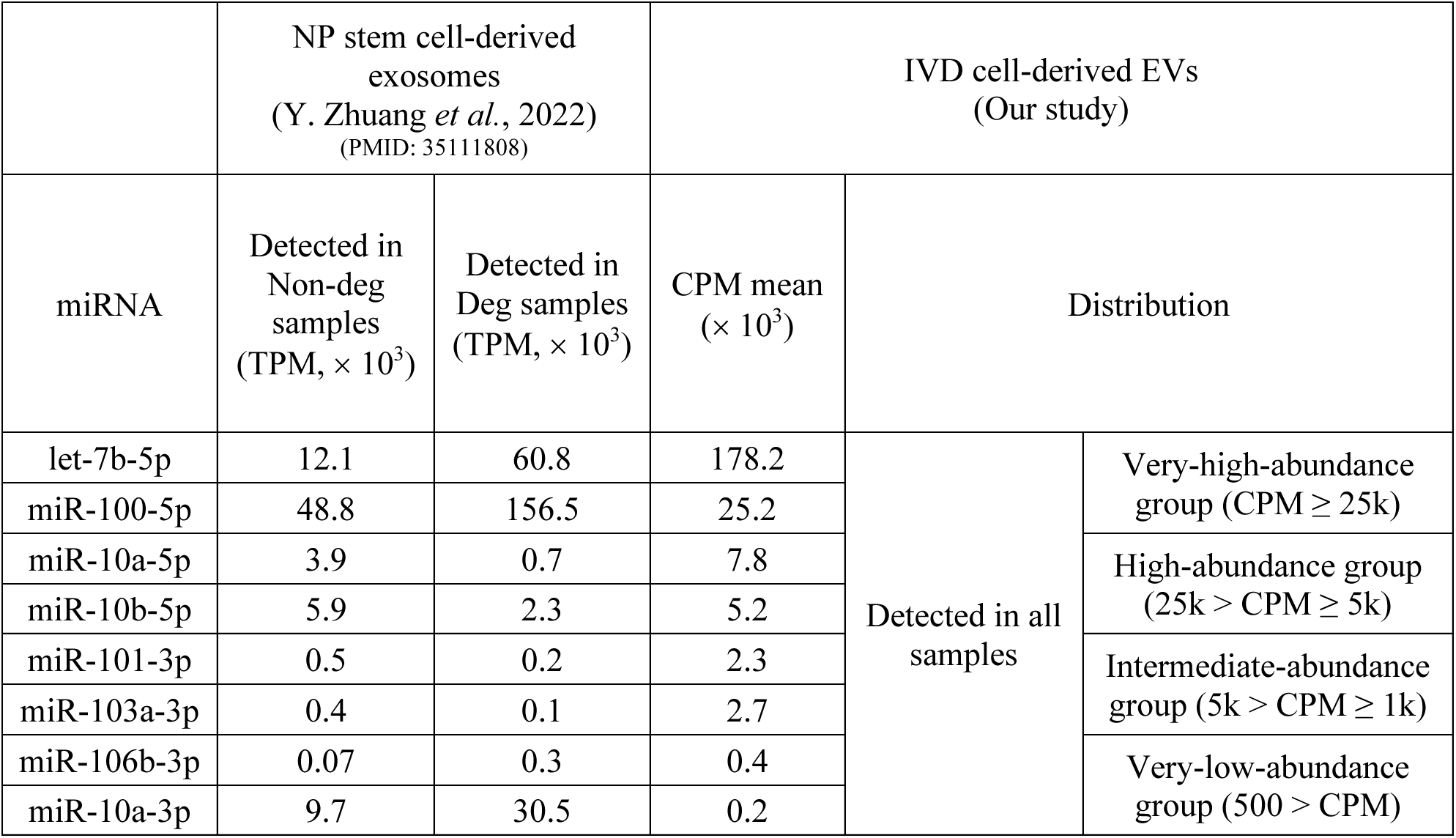
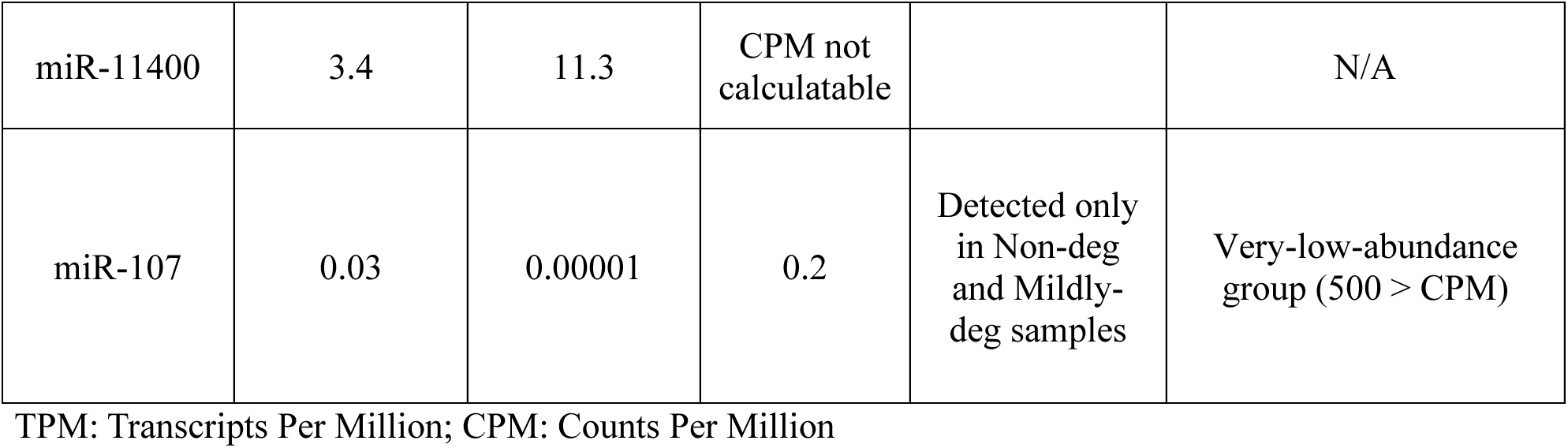
Comparison of human IVD cell-derived EV-enriched miRNAs and NP stem cell-derived exosome-enriched miRNAs.

NP cells and NPSC are both found within the NP region of IVDs. They share a common origin and exhibit both similar and distinct biological characteristics. NPSCs and NP cells both express MSC surface markers CD73, CD90, and CD105, but NPSCs express higher levels of these markers^67^, indicating a greater stemness. In addition, NPSCs lack the mature NP cell phenotypic marker CD24.^68^ These similarities and differences between NPSCs and NP cells may explain the specific miRNA profile and relative expression pattern in EVs derived from each cell source.

### 4.3 miRNA regulatory effects on cell phenotype and homeostasis in human IVD tissues

Previous studies showed a regulatory role in improving human IVD cell phenotypic markers, cellular homeostasis, and tissue integrity. It was reported that the relative expression of miR-129-5p was significantly reduced in NP from Deg compared with Non-deg tissue. Delivery of miR-129-5p via MSC-derived EVs improved cell viability, reduced cell apoptosis, and decreased ECM degradation in IL-1ß-induced NP cells.^69^ We detected miR-129-5p only in EVs derived from Non-deg samples. However, its relative expression was very low (CPM = 54). Similarly, a previous study showed that the relative expression of miR-499a-5p was reduced in NP from Deg compared with Non-deg tissues. Overexpression of miR-499a-5p decreased the levels of catabolic enzymes matrix metalloproteinase (MMP) 3 and 13 and increased the synthesis of ECM compositions type II collagen and aggrecan in TNF-α-induced NP cells.^70^ However, we detected a weak signal of miR-499a-5p in all EV samples, and its relative expression was too low to be calculable using the CPM method. In addition, Zhuang *et al.* reported that let-7b-5p inhibited cell proliferation, migration, and matrix synthesis while promoting apoptosis in human AF cells isolated from Deg IVD tissue.^66^ We did not detect any significant regulatory effect of let-7b-5p on the cell phenotypic and homeostatic markers. The differences between our results and those of previous studies indicate that selective packaging systems may exist in human IVD cells. IVD tissue-enriched miRNAs with therapeutic potentials are not necessarily selected and packaged into IVD cell-derived EVs for cell communication, cargo transportation, and/or signalling transduction.

We observed that the EV-enriched miR-100-5p demonstrated an upregulatory effect on *TIE2* under non-inflammatory conditions. When combined with let-7b-5p, the combination treatment upregulated *TIE2* and *FOXF1* under non-inflammatory conditions and *TIE2* under inflammatory conditions. These findings align with prior reports that miR-100-5p supports cellular homeostasis by engaging growth and survival pathways: it promotes myoblast proliferation while restraining differentiation via Trib2/mTOR/S6K signalling^71^ and, by targeting homeodomain-interacting protein kinase 2, it activates PI3K/AKT signalling in endothelial cells.^72^

### 4.4 Regulation of inflammatory response of miRNAs on human IVD and other musculoskeletal cells

MiRNAs play pivotal roles in regulating inflammatory responses in IVD and other musculoskeletal tissues, with effects varying by miRNA type and cellular context. For instance, Cao *et al.* reported that inhibition of let-7b-5p reduced pro-inflammatory mediators like *IL-6*, *IL-8*, and *MMP10* in chondrocytes, suggesting an inflammatory role.^73^ However, Palamà *et al.* found no significant change in *IL-6*, *IL-8*, or *COX-2* levels with let-7b-5p mimic treatment in IL-1ß-induced human articular chondrocytes.^74^ In our study, the let-7b-5p mimic significantly downregulated the pro-inflammatory mediators *IL-1ß* at the gene expression level and IL-6, IL-1α, TNF-α, CXCL10, and BDNF at the protein level. In addition, the let-7b-5p mimic significantly upregulated the anti-inflammatory mediators CCL22 and IL-10 at the protein level. Our findings indicate an anti-inflammatory role of let-7b-5p. These conflicting findings suggest that let-7b-5p’s effects may depend on cell type and microenvironment, warranting further investigation.

Similarly, miR-100-5p has demonstrated anti-inflammatory properties. It was shown to inhibit reactive oxygen species production and apoptosis in human chondrocytes^75^, while Gao *et al.* reported that exosomal miR-100-5p from human umbilical cord MSCs attenuated TNF-α, IL-1ß, IL-6, and IL-8 in mouse eosinophils.^76^ In our study, the miR-100-5p mimic significantly downregulated *IL-1ß* gene expression level and IL-6, IL-1α, TNF-α, CXCL10, BDNF, alongside a significant upregulation of CCL22 and IL-10 at the protein level, further supporting its anti-inflammatory property.

The combination treatment of let-7b-5p and miR-100-5p significantly downregulated 10 out of 12 pro-inflammatory mediators and significantly upregulated 2 out of 3 anti-inflammatory mediators, indicating strong anti-inflammatory properties. The expression of IL-1ß, IL-8, CXCL1, CCL24, and IFN-γ was significantly downregulated by the combination treatment, but not by the single-miRNA treatment. No statistically significant difference was observed in the multiple comparisons between the three treatment groups, which suggests there was no added effect from the combination compared with the single-miRNA treatment. More investigations are needed to further evaluate the potential of the added effects of combination treatment.

### 4.5 Regulation of cellular senescence of miRNAs on human IVD and other musculoskeletal cells

The regulation of cellular senescence of let-7b-5p is multifaceted. It was shown that let-7b-5p decreased the expression of p53 at the gene and protein expression levels in the human acute myeloid leukemia cells (K562 and HL-60), which was reversed when a let-7b-5p inhibitor was used.^77^ In contrast, Cao *et al.* demonstrated that inhibiting let-7b-5p reversed the increased cellular senescence in chondrocytes, indicating a pro-senescence role of let-7b-5p.^73^ Our study showed that let-7b-5p or the combination of let-7b-5p and miR-100-5p significantly downregulated the expression of the cyclin-dependent kinase inhibitor *p16*, which is strongly associated with reducing cellular senescence and the progression of tissue degeneration in IVDs. We did not detect significant dysregulation of *p53* under single-miRNA or combination treatment. The function of let-7b-5p, appears to be context-dependent, in addition, although miRNA inhibitors are designed to specifically bind and silence a specific miRNA, off-target effects have been reported that could explain the differences. More investigations are required to determine the regulatory roles of let-7b-5p and miR-100-5p on cellular senescence in human IVD cells, as such miRNAs could be particularly valuable in cell-free therapies aimed at managing cellular aging processes in IVD degeneration and other degenerative musculoskeletal diseases.

Taken together, miR-100-5p and let-7b-5p displayed considerable ECM preservation properties, anti-senescence, and anti-inflammatory effects in IVD cells by modulating transcriptional regulation and protein production. Their distinct yet complementary roles underscore the therapeutic potential of targeting these miRNAs to preserve cell phenotype, cellular homeostasis, and decrease inflammatory microenvironment and cellular senescence in IVD degeneration. Our study demonstrates the potential of EV-enriched miRNAs to modulate degenerative pathways in human IVD cells. Our findings open avenues for developing EV-enriched miRNA-based, cell-free therapies targeting IVD degeneration and related disorders. By focusing on miRNAs like miR-100-5p and let-7b-5p, our study not only elucidates the biological roles of these molecules in degenerative diseases but also supports further research into the combinatorial use of miRNAs for therapeutic applications from the EV perspective.

## 5 Conclusion

The selective packaging of miRNAs in IVD cell-derived EVs, particularly miR-100-5p and let-7b-5p, illustrates their relevance in modulating key cellular pathways. Our findings validated their functions in improving IVD cell phenotypic marker expression and homeostasis, counteracting the inflammatory microenvironment, and potentially reducing cellular senescence. Our study highlights the critical role of EV-enriched miRNAs in these aspects. EV-enriched miRNAs hold promise for developing cell-free regenerative therapies, particularly for treating IVD degeneration and potentially other musculoskeletal disorders. Further exploration of miRNA combinatorial therapies may provide novel insights into enhancing therapeutic efficacy and specificity.

## Author contributions

Conceptualization L.L. and L.H.; Methodology, L.L. and H.A.; Validation, L.L., A.S., S.G., H.C. and L.H.; Formal analysis, L.L.; Investigation, L.L.; Resources, K.U., S.A., J.O., P.J. and L.H.; Data curation, L.L. and A.S.; Writing—original draft preparation, L.L.; Writing—review and editing, L.L., A.S., S.G., H.A., K.U., S.A., J.O., P.J., H.C. and L.H.; Visualization, L.L.; Supervision, H.C. and L.H.; Project administration, L.H.; Funding acquisition, L.H. All authors have read and agreed to the current version of the manuscript.

## Data availability statement

All data generated or analyzed during this study are included in the manuscript and supporting files. The raw data and materials used to support the findings of this study are available from the corresponding author upon request.

## Patient consent statement

Informed consent was obtained from all subjects involved in the study.

## Ethics approval statement

The study was conducted in accordance with the Declaration of Helsinki and approved by the Institutional Review Board of McGill University (IRB #00010120).

## Acknowledgments

The authors would like to acknowledge the Institut de recherche en immunologie et en cancérologie, and staff (Raphaëlle Lambert and Patrick Gendron) for services provided. This work was supported by the Canadian Institutes of Health Research under grant #PJT-178111; Le Réseau de Recherche en Santé Buccodentaire et Osseuse major infrastructure grant; ThéCell and Le Fonds de Recherche du Québec-Santé doctoral training award (#272079) to Li Li.

