## Supplementary materials for "Beyond miRNA cargo profiles: anti-inflammatory roles of extracellular vesicle-enriched miRNAs derived from human intervertebral disc cells unveiled by functional testing"

### 1. Supplementary Methods

### 2. Supplementary Results

2.1 Comparisons of the pathways and functional annotations of the shared miRNAs detected in two of the three groups.

### 3. Supplementary Figures and Tables

**Figure S1**. Comparisons of the top 10 musculoskeletal disease-associated annotations of the unique miRNAs exclusively detected in the non-degenerate, mildly-degenerate, and degenerate samples.

**Figure S2**. The top 10 musculoskeletal disease-associated annotations of miRNAs, exclusively detected in two of the three groups.

**Figure S3**. The top 10 musculoskeletal disease-associated pathways and annotations of the shared miRNAs by abundance.

**Figure S4**. Toll-like receptor (TLR) activation and the regulatory effects of miRNA mimic combination treatments on inflammatory mediators and senescence markers in non-inflammatory condition.

**Figure S5**. TLR activation at the protein level.

**Table S1**. Shared and unique top 10 musculoskeletal disease-associated GO-BP annotations of unique miRNAs detected in the non-degenerate, mildly-degenerate, and degenerate samples.

**Table S2**. Shared and unique top 10 musculoskeletal disease-associated GO-CC annotations of unique miRNAs detected in the non-degenerate, mildly-degenerate, and degenerate samples.

**Table S3**. Shared and unique top 10 musculoskeletal disease-associated GO-MF annotations of unique miRNAs detected in the non-degenerate, mildly-degenerate, and degenerate samples.

**Table S4**. Shared and unique top 10 musculoskeletal disease-associated GO-BP annotations of miRNAs detected in two of the Non-deg, Mildly-deg, and Deg groups.

**Table S5**. Shared and unique top 10 musculoskeletal disease-associated GO-CC annotations of miRNAs detected in two of the Non-deg, Mildly-deg, and Deg groups.

**Table S6**. Shared and unique top 10 musculoskeletal disease-associated GO-MF annotations of miRNAs detected in two of the Non-deg, Mildly-deg, and Deg groups.

**Table S7**. The musculoskeletal disease-associated pathways and annotations of the shared miR-148a-5p and miR-210-3p (CPM > 300) in the non-degenerate and mildly-degenerate samples.

**Table S8**. The musculoskeletal disease-associated KEGG pathways of the shared miRNAs with CPM > 300 in the non-degenerate and degenerate samples.

**Table S9**. The musculoskeletal disease-associated GO-BP annotations of the shared miRNAs with CPM > 300 in the non-degenerate and degenerate samples.

**Table S10**. The musculoskeletal disease-associated GO-CC annotations of the shared miRNAs with CPM > 300 in the non-degenerate and degenerate samples.

**Table S11**. The musculoskeletal disease-associated GO-MF annotations of the shared miRNAs with CPM > 300 in the non-degenerate and degenerate samples.

**Table S12**. The musculoskeletal disease-associated GO-CC annotations of the shared miR-10b-3p (CPM > 300) in the mildly-degenerate and degenerate samples.

**Table S13**. Concentration of inflammatory mediators.

### 1. Supplementary methods

***Evaluation metrics***: We set five criteria for pathway and annotation selection. 1) Relevance to musculoskeletal tissue types. We focused on pathways and annotations directly impacting intervertebral discs (IVDs), bones, joints, muscles, tendons, ligaments, and connective tissues. Pathways that influence cellular processes within these tissues, such as ECM remodelling, bone formation, and muscle contraction, were prioritized. 2) Association with inflammatory and degenerative diseases. Many musculoskeletal conditions, such as IVD degeneration, osteoarthritis, rheumatoid arthritis, and musculoskeletal pain syndromes, are driven by inflammation and tissue degeneration. Therefore, pathways linked to inflammatory mediators, ECM degradation, and cartilage/bone remodelling were prioritized. 3) Pathways involved in repair and regeneration. Pathways that support tissue healing, such as those involved in stem cell activity and muscle regeneration, are considered highly relevant to musculoskeletal health, and were prioritized. 4) Evidence from musculoskeletal disease studies. We considered the extent to which each pathway was mentioned in the literature specifically related to musculoskeletal diseases. Pathways frequently studied or linked to biomarkers of musculoskeletal diseases were given higher priority. 5) Priority of musculoskeletal disease association in functional overlaps. Some pathways share functional roles, particularly those related to cell signalling (e.g., integrin signalling, Wnt signalling). When functional overlap was present, the pathways with the most direct or documented impact on musculoskeletal conditions were prioritized. We used databases such as Kyoto Encyclopedia of Genes and Genomes (KEGG), PubMed, and NCBI to search for pathways and keywords relevant to musculoskeletal diseases.

***Algorithmic process***: 1) Initial screening: each pathway/annotation from the list exported in the function prediction step was first reviewed to assess its basic function (e.g., cell cycle, apoptosis) and any documented relevance to musculoskeletal diseases. 2) Relevance scoring: pathways and annotations were assigned a relevance score (on a scale of 1-10) based on the five criteria described above. This score was derived from keyword matches (e.g., IVD, bone, and muscle) with relevant biological processes or diseases. 3) Clustering of similar pathways/annotations: pathways and annotations with similar biological roles (e.g., actin binding and cytoskeletal protein binding) were grouped to avoid redundancy. The pathway/annotation with the highest score in each group was retained. 4) Top pathway/annotation selection: the top 10 pathways/annotations were chosen based on the highest relevance scores, ensuring they covered a range of functions important for musculoskeletal biology (e.g., inflammation, tissue regeneration, structural integrity). The selected pathways/annotations were ranked by merged FDR and were presented as Pathways union vs. -$\log_{10}$ (Merged FDR) in histograms. When the merged FDR = 0, the -$\log_{10}$ (Merged FDR) was presented as infinity ($\infty$).

### 2. Supplementary results

#### *2.1 Comparisons of the pathways and functional annotations of the shared miRNAs detected in two of the three groups*

In the Non-deg and Mildly-deg groups, enrichment of cell cycle and p53 signalling pathway indicates early shifts toward checkpoint activation and senescence from a healthy baseline, while Hippo and TGF-beta signalling pathways reflect re-tuning of mechanosensing and matrix-repair programs. Ubiquitin-mediated proteolysis points to proteostasis and inflammatory signal turnover. Together with the shared pathways, this profile depicts an incipient transition in growth control, matrix remodelling, and stress handling that may still be tractable to intervention. In the Non-deg and Deg groups, the pairing retains growth-control and repair axes, such as cell cycle, p53, Hippo, TGF-beta, ubiquitin proteolysis signalling pathways, but is distinguished by regulation of the actin cytoskeleton, consistent with degeneration-anchored cytoskeletal and adhesion reprogramming that amplifies inflammatory and catabolic outputs. The presence of AMPK signalling pathway suggests metabolic stress adaptation and pro-autophagy signalling that restrains NF-κB activity, while adherens junction highlights junctional integrity as a determinant of survival and mechano-signalling. This constellation links energy balance, cell-cell cohesion, and cytoskeletal mechanics to the healthy-to-degenerate transition. In the Mildly-deg and Deg group, enrichment of FoxO, MAPK, and Wnt signalling pathways indicates stress-response and autophagy programs, cytokine and mechanical-stress signalling driving MMP/ADAMTS, and phenotype and matrix-turnover control. Convergence on actin cytoskeleton further supports altered stiffness sensing and traction forces in degeneration. Unique enrichment of apoptosis, tight junction, vascular endothelial growth factor (VEGF), and mechanistic target of rapamycin (mTOR) signalling pathways aligns with latestage features, such as cell loss, barrier and polarity disruption, pathological neovascularization and nerve ingrowth, and suppressed autophagy with metabolic dysregulation, collectively explaining reduced matrix maintenance and heightened pain-associated signalling.

Gene ontology (GO)- biological process (BP) (**Table S4**), -cellular component (CC) (**Table S5**), and -molecular function (MF) (**Table S6**) converge on growth-control and mechano-repair programs in the Non-deg and Mildly-deg groups: cell cycle/division, Hippo and Wnt signalling, autophagy/ubiquitin-proteasome, and actin-cytoskeleton reorganization, localized to extracellular exosomes, focal adhesions, adherens junctions, and cytoskeletal/nucleolar compartments, with cadherin/integrin/SMAD/vinculin interactions. Together, these features indicate a pro-survival, mechanoresponsive state with active proteostasis and EV-mediated communication that support matrix maintenance. We identified miR-148a-5p and miR-210-3p with CPM > 300. They are associated with growth-control pathways and are representative of the functional prediction of miRNAs shared by the Non-deg and Mildly-deg groups (**Table S7**). Annotations emphasize ECM and adhesion remodelling alongside developmental and growth-factor cues in the Non-deg and Deg groups: ECM organization, osteoblast/BMP and Wnt pathways, and actin cytoskeleton organization (GO-BP) (**Table S4**); focal/adherens junctions, laminin/basement-membrane and integrin complexes (GO-CC) (**Table S5**); and cadherin/SMAD binding with transcription-factor and ubiquitin-transferase activities (GO-MF) (**Table S6**). This profile reflects the healthy-to-degenerate transition, linking matrix restructuring and junctional mechanics to gene-regulatory control. We identified miR-21-3p, miR-374b-5p, miR-130a-3p, miR-485-3p, and miR-135b-5p with CPM > 300 in the Non-deg and Deg group pair. The expression of miR-21-3p is higher in the Non-deg group, while that of others is higher in the Deg group. In particular, miR-21-3p is associated with ECM and adhesion remodelling pathways. MiR-374b-5p and miR-130a-3p are associated with cellular senescence- and angiogenesis-associated pathways and annotations (**Table S8-S11**). Terms cluster around nuclear/chromatin and kinase-driven control with stress/survival balance in the Mildly-deg and Deg groups: chromatin organization, cell-cycle regulation, protein phosphorylation, apoptosis and PI3K signalling (GO-BP) (**Table S4**); nucleoplasm/nuclear matrix/mTOC and spindle assemblies with residual collagen compartments (GO-CC) (**Table S5**); and chromatin/transcription factor interfaces, serine/threonine-kinase and ubiquitin-transferase activities (GO-MF) (**Table S6**), with persistent cadherin binding. Collectively, this indicates entrenched transcriptional reprogramming and signalling dependency in mid-to-late degeneration. MiR-10b-3p was the only miRNA with CPM > 300 detected in the Mildly-deg and Deg group pair. It was associated with annotations only in GO-CC term, which are nucleoplasm, nucleus, and cytosol (**Table S12**).

#### 3. Supplementary figures and tables

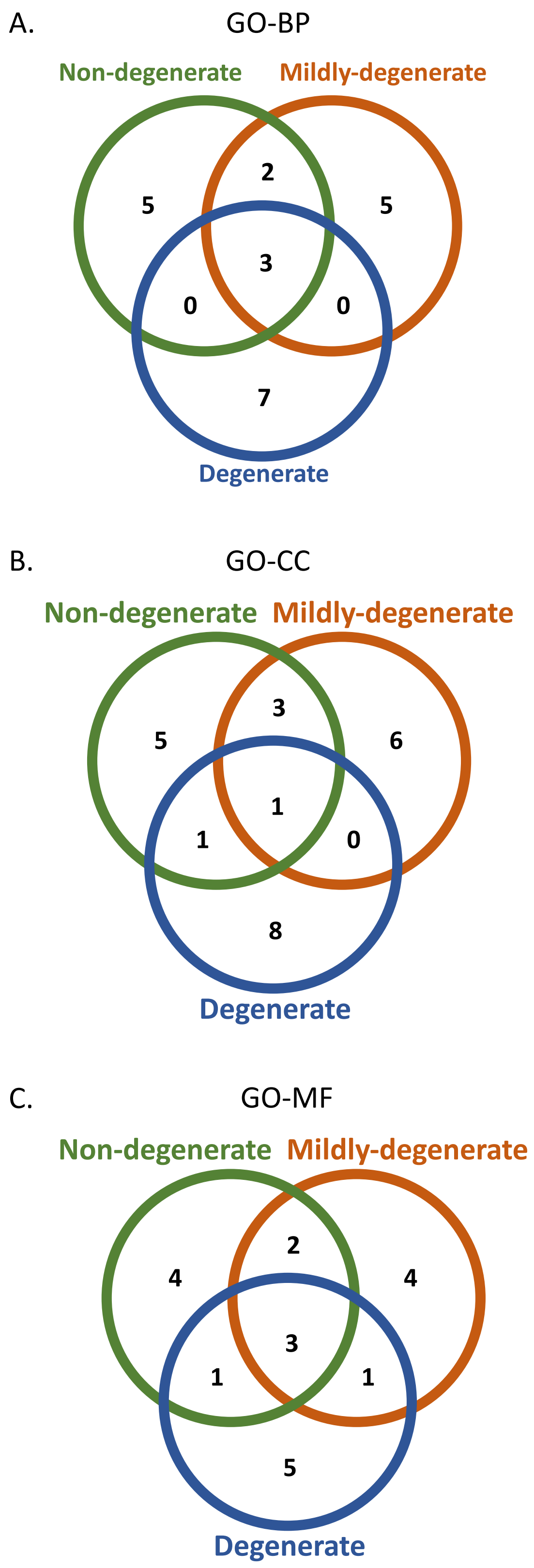

**Figure S1. Comparisons of the top 10 musculoskeletal disease-associated annotations of the unique miRNAs exclusively detected in the Non-deg, Mildly-deg, and Deg samples.** Venn diagrams presenting the top 10 overlapping and unique musculoskeletal disease-associated Gene Ontology (GO) annotations within the (**A**) biological process (BP), (**B**) cellular component (CC), and (**C**) molecular function (MF) terms with strong association with EV cargo miRNAs exclusively detected in the non-degenerate (Non-deg), mildly-degenerate (Mildly-deg), and degenerate (Deg) samples. Non-deg (green), Mildly-deg (orange), and Deg (blue).

**Table S1. Shared and unique top 10 musculoskeletal disease-associated GO-BP annotations of miRNAs detected in the Non-deg, Mildly-deg, and Deg samples.**

| Groups | Quantity of shared pathways | Pathways |
|---|---|---|
| Detected in the Non-deg, Mildly-deg, and Deg groups | 3 | Cell cycle |
| | | Cell division |
| | | Protein phosphorylation |
| Detected in the Non-deg and Mildly-deg groups | 2 | TGF-beta receptor signalling pathway |
| | | Apoptotic process |
| Exclusively detected in the Non-deg group | 5 | Regulation of cell migration |
| | | Wnt signalling pathway |
| | | Cytoskeleton organization |
| | | Osteoblast differentiation |
| | | VEGF receptor signalling pathway |
| Exclusively detected in the Mildly-deg group | 5 | Cell-matrix adhesion |
| | | Cellular response to DNA damage stimulus |
| | | Phosphorylation |
| | | Positive regulation of osteoblast differentiation |
| | | Regulation of osteoblast differentiation |
| **Exclusively detected in the Deg group** | **7** | Collagen fibril organization |
| | | Chromatin organization |
| | | Response to oxidative stress |
| | | Adherens junction assembly |
| | | Protein polyubiquitination |
| | | Cellular response to growth factor stimulus |
| | | Transmembrane receptor protein serine/threonine kinase signalling pathway |

**Table S2. Shared and unique top 10 musculoskeletal disease-associated GO-CC annotations of unique miRNAs detected in the Non-deg, Mildly-deg, and Deg samples.**

| Groups | Quantity of shared pathways | Pathways |
|---|---|---|
| Detected in the Non-deg, Mildly-deg, and Deg groups | 1 | Focal adhesion |
| Detected in the Non-deg and Mildly-deg groups | 3 | Cytoskeleton |
| | | Cell-cell junction |
| | | Adherens junction |
| Detected in the Non-deg and Deg groups | 1 | Collagen-containing extracellular matrix |
| Exclusively detected in the Non-deg group | 5 | Laminin complex |
| | | Basement membrane |
| | | Actin cytoskeleton |
| | | Sarcolemma |
| | | Microtubule cytoskeleton |
| Exclusively detected in the Mildly-deg group | 6 | Cell-substrate junction |
| | | Microtubule |
| | | Ubiquitin ligase complex |
| | | Integrin alpha3-beta1 complex |
| | | RISC complex |
| | | SMAD protein complex |
| Exclusively detected in the Deg group | 8 | Cell junction |
| | | Stress fiber |
| | | Dendritic spine |
| | | Neuron projection |
| | | Costamere |
| | | Extracellular matrix |
| | | Actin cap |
| | | Collagen type I trimer |

**Table S3. Shared and unique top 10 musculoskeletal disease-associated GO-MF annotations of unique miRNAs detected in the Non-deg, Mildly-deg, and Deg samples.**

| Groups | Quantity of shared pathways | Pathways |
|---|---|---|
| Detected in the Non-deg, Mildly-deg, and Deg groups | 3 | Protein kinase activity |
| | | SMAD binding |
| | | Cadherin binding |
| Detected in the Non-deg and Mildly-deg groups | 2 | Ubiquitin-protein transferase activity |
| | | Histone binding |
| Detected in the Non-deg and Deg groups | 1 | Actin binding |
| Detected in the Mildly-deg and Deg groups | 1 | Protein serine/threonine kinase activity |
| Exclusively detected in the Non-deg group | 4 | Integrin binding |
| | | Metal ion binding |
| | | Cyclin-dependent protein serine/threonine kinase regulator activity |
| | | GTPase binding |
| Exclusively detected in the Mildly-deg group | 4 | Mitogen-activated protein kinase kinase binding |
| | | Proteoglycan binding |
| | | Laminin binding |
| | | Transcription factor binding |
| Exclusively detected in the Deg group | 5 | Extracellular matrix structural constituent conferring tensile strength |
| | | Ubiquitin-ubiquitin ligase activity |
| | | TGF-beta binding |
| | | TGF-beta-activated receptor activity |
| | | Beta-catenin binding |

**Table S4. Shared and unique top 10 musculoskeletal disease-associated GO-BP annotations of miRNAs detected in two of the Non-deg, Mildly-deg, and Deg groups.**

| Groups | Quantity of shared pathways | Pathways |
|---|---|---|
| Shared across Non-deg/Mildly-deg, Non-deg/Deg, and Mildly-deg/Deg | 3 | Cell cycle |
| | | Cell division |
| | | Wnt signalling pathway |
| Non-deg/Mildly-deg and Non-deg/Deg | 2 | Osteoblast differentiation |
| | | TGF-beta receptor signalling pathway |
| Non-deg/Mildly-deg | 5 | Autophagy |
| | | Hippo signalling |
| | | Proteasome-mediated ubiquitin-dependent protein catabolic process |
| | | Protein ubiquitination |
| | | Regulation of actin cytoskeleton reorganization |
| Non-deg/Deg | 5 | Actin cytoskeleton organization |
| | | BMP signalling pathway |
| | | Extracellular matrix organization |
| | | Fibroblast growth factor receptor signalling pathway |
| | | Skeletal system development |
| Mildly-deg/Deg | 7 | Apoptotic process |
| | | Chromatin organization |
| | | Phosphatidylinositol 3-kinase signalling |
| | | Protein phosphorylation |
| | | Regulation of cell cycle |
| | | Regulation of transcription by RNA polymerase II |
| | | Ubiquitin-dependent protein catabolic process |

**Table S5. Shared and unique top 10 musculoskeletal disease-associated GO-CC annotations of miRNAs detected in two of the Non-deg, Mildly-deg, and Deg groups.**

| Groups | Quantity of shared pathways | Pathways |
|---|---|---|
| Shared across Non-deg/Mildly-deg, Non-deg/Deg, and Mildly-deg/Deg | 4 | Actin cytoskeleton |
| | | Cytoskeleton |
| | | Focal adhesion |
| | | Spindle |
| Non-deg/Mildly-deg and Non-deg/Deg | 1 | Adherens junction |
| Non-deg/Deg and Mildly-deg/Deg | 1 | Lamellipodium |
| Non-deg/Mildly-deg | 5 | Collagen type I trimer |
| | | Endoplasmic reticulum |
| | | Extracellular exosome |
| | | Nucleolus |
| | | Ribonucleoprotein complex |
| Non-deg/Deg | 4 | Collagen-containing extracellular matrix |
| | | Extracellular matrix |
| | | Microtubule cytoskeleton |
| | | Nuclear envelope |
| Mildly-deg/Deg | 5 | Chromatin |
| | | Collagen trimer |
| | | Microtubule organizing center |
| | | Nuclear matrix |
| | | Ubiquitin ligase complex |

**Table S6. Shared and unique top 10 musculoskeletal disease-associated GO-MF annotations of miRNAs detected in two of the Non-deg, Mildly-deg, and Deg groups.**

| Groups | Quantity of shared pathways | Pathways |
|---|---|---|
| Shared across Non-deg/Mildly-deg, Non-deg/Deg, and Mildly-deg/Deg | 1 | Cadherin binding |
| Non-deg/Mildly-deg and Non-deg/Deg | 3 | Actin binding |
| | | Integrin binding |
| | | SMAD binding |
| Non-deg/Deg and Mildly-deg/Deg | 3 | Protein serine/threonine kinase activity |
| | | Transcription factor binding |
| | | Ubiquitin-protein transferase activity |
| Non-deg/Mildly-deg | 6 | Ephrin receptor binding |
| | | GTPase binding |
| | | Platelet-derived growth factor binding |
| | | Structural constituent of cytoskeleton |
| | | TGF beta-activated receptor activity |
| | | Vinculin binding |
| Non-deg/Deg | 3 | DNA-binding transcription factor activity |
| | | Extracellular matrix structural constituent |
| | | Nucleosome binding |
| Mildly-deg/Deg | 6 | ATP binding |
| | | Chromatin binding |
| | | Histone binding |
| | | Protein kinase activity |
| | | Protein-containing complex binding |
| | | RNA polymerase II cis-regulatory region sequence-specific DNA binding |

**Table S7. The musculoskeletal disease-associated pathways and annotations of the shared miR-148a-5p and miR-210-3p (CPM > 300) in the Non-deg and Mildly-deg samples.**

| | KEGG | | GO-BP | | GO-CC | | GO-MF | |
|---|---|---|---|---|---|---|---|---|
| | miR-148a-5p | miR-210-3p | miR-148a-5p | miR-210-3p | miR-148a-5p | miR-210-3p | miR-148a-5p | miR-210-3p |
| 1 | Cell cycle | No significant results | Protein phosphorylation | No significant results | Cytoplasm | Cytoplasm | Protein binding | Protein binding |
| 2 | FoxO signalling pathway | | Regulation of cell cycle | | Cytosol | Nucleoplasm | RNA binding | Ubiquitin-protein transferase activity |
| 3 | | | Wnt signalling pathway, calcium modulating pathway | | Nucleus | Cytosol | ATP binding | Enzyme binding |
| 4 | | | | | Nucleoplasm | Nucleus | | Ubiquitin protein ligase activity |
| 5 | | | | | P-body | | | |
| 6 | | | | | Nucleolus | | | |
| 7 | | | | | RISC complex | | | |
| 8 | | | | | Centrosome | | | |
| 9 | | | | | Beta-catenin-TCF complex | | | |
| 10 | | | | | Extracellular exosomes | | | |

**Table S8. The musculoskeletal disease-associated KEGG pathways of the shared miRNAs with CPM > 300 in the Non-deg and Deg samples.**

| | miR-21-3p | miR-374b-5p | miR-130a-3p | miR-485-3p | miR-135b-5p |
|---|---|---|---|---|---|
| 1 | Proteoglycans in cancer | p53 signalling pathway | Autophagy - animal | No significant results | No significant results |
| 2 | Cell cycle | FoxO signalling pathway | mTOR signalling pathway | | |
| 3 | Protein processing in endoplasmic reticulum | Cell cycle | FoxO signalling pathway | | |
| 4 | Regulation of actin cytoskeleton | TGF-beta signalling pathway | TGF-beta signalling pathway | | |
| 5 | Fluid shear stress and atherosclerosis | | Cell cycle | | |
| 6 | Ubiquitin-mediated proteolysis | | AMPK signalling pathway | | |
| 7 | AMPK signalling pathway | | PI3K-Akt signalling pathway | | |
| 8 | Autophagy - animal | | Apoptosis | | |
| 9 | MAPK signalling pathway | | Focal adhesion | | |
| 10 | Focal adhesion | | Ubiquitin-mediated proteolysis | | |

**Table S9. The musculoskeletal disease-associated GO-BP annotations of the shared miRNAs with CPM > 300 in the Non-deg and Deg samples.**

| | miR-21-3p | miR-374b-5p | miR-130a-3p | miR-485-3p | miR-135b-5p |
|---|---|---|---|---|---|
| 1 | Cell migration | Negative regulation of transcription by RNA polymerase II | Cell cycle | No significant results | No significant results |
| 2 | Regulation of cell cycle | Cell cycle | Chromatin organization | | |
| 3 | Ubiquitin-dependent protein catabolic process | Cell division | Protein phosphorylation | | |
| 4 | Cell cycle | DNA damage response, signal transduction by p53 class mediator resulting in cell cycle arrest | Protein ubiquitination | | |
| 5 | VEGF receptor signalling pathway | Peptidyl-serine phosphorylation | Regulation of transcription by RNA polymerase II | | |
| 6 | Regulation of translation | Fibroblast growth factor receptor signalling pathway | Regulation of cell cycle | | |
| 7 | Apoptotic process | Regulation of translation | DNA damage response, signal transduction by p53 class mediator resulting in cell cycle arrest | | |
| 8 | Angiogenesis | Protein phosphorylation | Cell division | | |
| 9 | Wnt signalling pathway, calcium modulating pathway | Endoplasmic reticulum tubular network organization | Apoptotic process | | |
| 10 | Protein phosphorylation | Activation of MAPKK activity | Ubiquitin-dependent protein catabolic process | | |

**Table S10. The musculoskeletal disease-associated GO-CC annotations of the shared miRNAs with CPM > 300 in the Non-deg and Deg samples.**

| | miR-21-3p | miR-374b-5p | miR-130a-3p | miR-485-3p | miR-135b-5p |
|---|---|---|---|---|---|
| 1 | Nucleus | Nucleus | Cytoskeleton | Cytoplasm | Cytosol |
| 2 | Cytoplasm | Nucleoplasm | Golgi apparatus | Cytosol | Cytoplasm |
| 3 | Focal adhesion | Cytosol | Chromatin | Nucleus | Focal adhesion |
| 4 | Chromatin | Cytoplasm | Microtubule | Nucleoplasm | |
| 5 | Growth cone | Ubiquitin ligase complex | Focal adhesion | Cell junction | |
| 6 | Cytoskeleton | Nuclear envelope | Nuclear envelope | Cytoskeleton | |
| 7 | Lamellipodium | Ribonucleoprotein complex | Endoplasmic reticulum | Perinuclear region of cytoplasm | |
| 8 | Actin cytoskeleton | Endoplasmic reticulum | Actin cytoskeleton | Axon | |
| 9 | Centrosome | Endoplasmic reticulum membrane | Adherens junction | Microtubule cytoskeleton | |
| 10 | Spindle | Protein-containing complex | Extracellular exosomes | Extracellular exosomes | |

**Table S11. The musculoskeletal disease-associated GO-MF annotations of the shared miRNAs with CPM > 300 in the Non-deg and Deg samples.**

| | miR-21-3p | miR-374b-5p | miR-130a-3p | miR-485-3p | miR-135b-5p |
|---|---|---|---|---|---|
| 1 | RNA binding | RNA binding | Protein binding | Protein binding | Protein binding |
| 2 | Protein binding | Protein binding | Cadherin binding | GTPase activity | Chromatin binding |
| 3 | Cadherin binding | Metal ion binding | Chromatin binding | | Nucleic acid binding |
| 4 | Protein kinase binding | DNA binding | Protein kinase activity | | Metal ion binding |
| 5 | ATP binding | Ubiquitin-protein transferase activity | Ubiquitin-protein transferase activity | | |
| 6 | SMAD binding | Transcription factor binding | DNA-binding transcription factor activity | | |
| 7 | Transcription factor binding | Nucleotide binding | Transcription factor binding | | |
| 8 | Chromatin binding | Protein serine/threonine kinase activity | Histone methyltransferase activity (h3-k4 specific) | | |
| 9 | Protein kinase activity | Histone deacetylase binding | Protein serine/threonine kinase activity | | |
| 10 | GTPase activator activity | DNA-binding transcription factor activity | Nucleosome binding | | |

**Table S12. The musculoskeletal disease-associated GO-CC annotations of the shared miR-10b-3p (CPM > 300) in the Mildly-deg and Deg samples.**

| | miR-10b-3p |
|---|---|
| 1 | nucleoplasm |
| 2 | nucleus |
| 3 | cytosol |

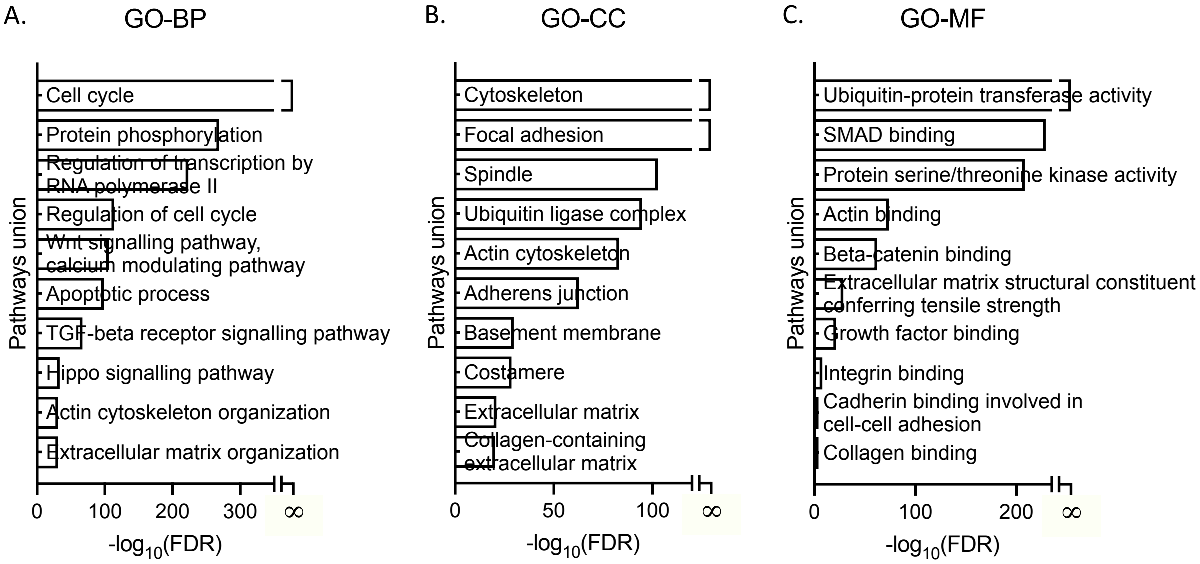


**Figure S2. The top 10 musculoskeletal disease-associated annotations of the shared 184 miRNAs detected in the Non-deg, Mildly-deg, and Deg samples.** The GO within the (**A**) BP, (**B**) CC, and (**C**) MF terms analyses presenting the top 10 musculoskeletal disease-associated annotations with strong association with the 184 EV cargo miRNAs detected in the Non-deg, Mildly-deg, and Deg samples. GO *p*-value threshold was set at 0.05. The x-axis represents the logarithmic scale of the FDR (-Log$_{10}$ (FDR)), and the y-axis describes the pathways union of miRNAs involved in the enriched GO annotations. When the merged FDR = 0, the -log$_{10}$ (Merged FDR) was presented as infinity (∞).

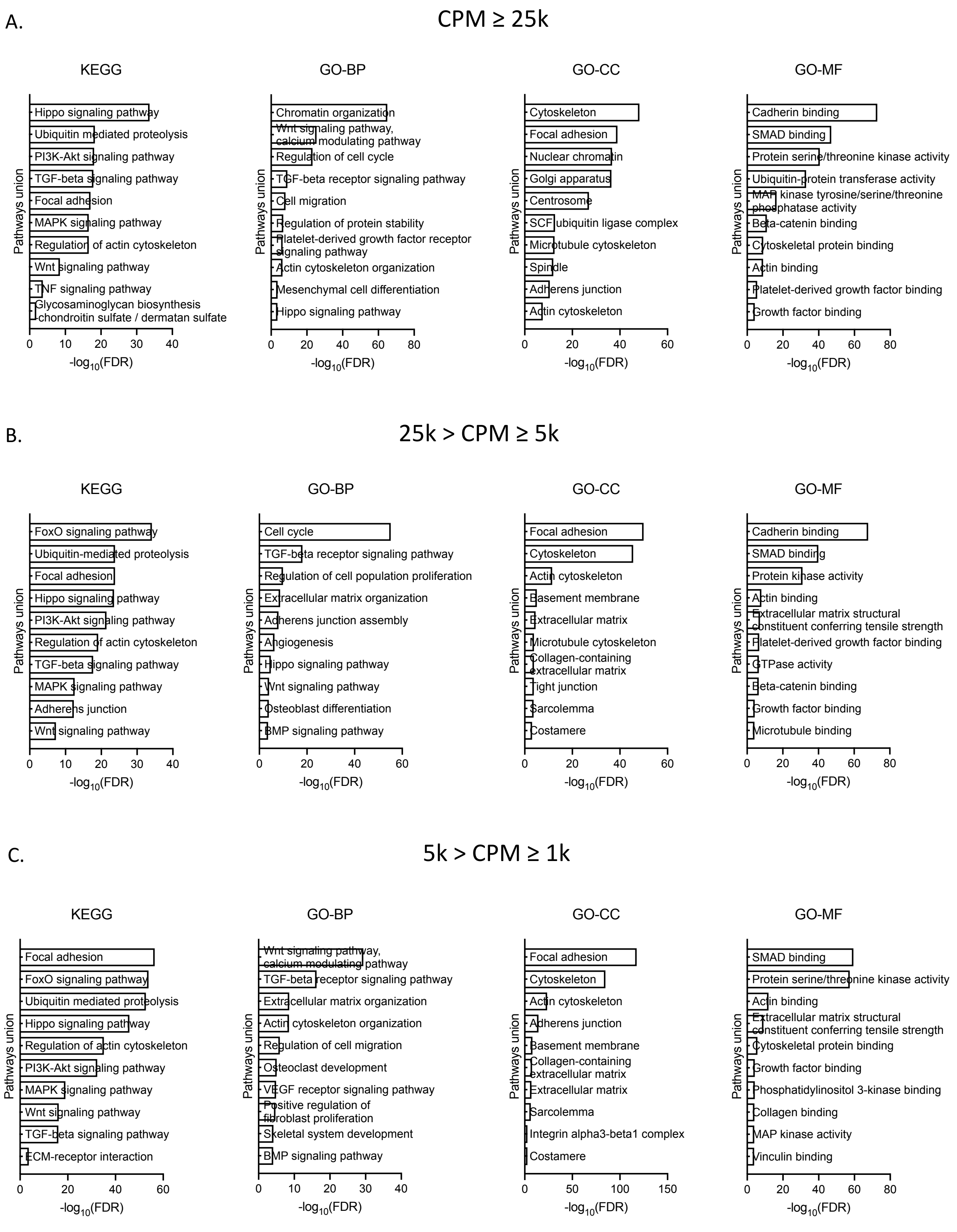
A.
CPM ≥ 25k
KEGG
Hippo signaling pathway
Ubiquitin mediated proteolysis
PI3K-Akt signaling pathway
TGF-beta signaling pathway
Focal adhesion
MAPK signaling pathway
Regulation of actin cytoskeleton
Wnt signaling pathway
TNF signaling pathway
Glycosaminoglycan biosynthesis chondroitin sulfate / dermatan sulfate
GO-BP
Chromatin organization
Wnt signaling pathway, calcium modulating pathway
Regulation of cell cycle
TGF-beta receptor signaling pathway
Cell migration
Regulation of protein stability
Platelet-derived growth factor receptor signaling pathway
Actin cytoskeleton organization
Mesenchymal cell differentiation
Hippo signaling pathway
GO-CC
Cytoskeleton
Focal adhesion
Nuclear chromatin
Golgi apparatus
Centrosome
SCF ubiquitin ligase complex
Microtubule cytoskeleton
Spindle
Adherens junction
Actin cytoskeleton
GO-MF
Cadherin binding
SMAD binding
Protein serine/threonine kinase activity
Ubiquitin-protein transferase activity
MAP kinase tyrosine/serine/threonine phosphatase activity
Beta-catenin binding
Cytoskeletal protein binding
Actin binding
Platelet-derived growth factor binding
Growth factor binding
Pathways union
-log10(FDR)
B.
25k > CPM ≥ 5k
KEGG
FoxO signaling pathway
Ubiquitin-mediated proteolysis
Focal adhesion
Hippo signaling pathway
PI3K-Akt signaling pathway
Regulation of actin cytoskeleton
TGF-beta signaling pathway
MAPK signaling pathway
Adherens junction
Wnt signaling pathway
GO-BP
Cell cycle
TGF-beta receptor signaling pathway
Regulation of cell population proliferation
Extracellular matrix organization
Adherens junction assembly
Angiogenesis
Hippo signaling pathway
Wnt signaling pathway
Osteoblast differentiation
BMP signaling pathway
GO-CC
Focal adhesion
Cytoskeleton
Actin cytoskeleton
Basement membrane
Extracellular matrix
Microtubule cytoskeleton
Collagen-containing extracellular matrix
Tight junction
Sarcolemma
Costamere
GO-MF
Cadherin binding
SMAD binding
Protein kinase activity
Actin binding
Extracellular matrix structural constituent conferring tensile strength
Platelet-derived growth factor binding
GTPase activity
Beta-catenin binding
Growth factor binding
Microtubule binding
Pathways union
-log10(FDR)
C.
5k > CPM ≥ 1k
KEGG
Focal adhesion
FoxO signaling pathway
Ubiquitin mediated proteolysis
Hippo signaling pathway
Regulation of actin cytoskeleton
PI3K-Akt signaling pathway
MAPK signaling pathway
Wnt signaling pathway
TGF-beta signaling pathway
ECM-receptor interaction
GO-BP
Wnt signaling pathway, calcium modulating pathway
TGF-beta receptor signaling pathway
Extracellular matrix organization
Actin cytoskeleton organization
Regulation of cell migration
Osteoclast development
VEGF receptor signaling pathway
Positive regulation of fibroblast proliferation
Skeletal system development
BMP signaling pathway
GO-CC
Focal adhesion
Cytoskeleton
Actin cytoskeleton
Adherens junction
Basement membrane
Collagen-containing extracellular matrix
Extracellular matrix
Sarcolemma
Integrin alpha3-beta1 complex
Costamere
GO-MF
SMAD binding
Protein serine/threonine kinase activity
Actin binding
Extracellular matrix structural constituent conferring tensile strength
Cytoskeletal protein binding
Growth factor binding
Phosphatidylinositol 3-kinase binding
Collagen binding
MAP kinase activity
Vinculin binding
Pathways union
-log10(FDR)

D. 1k > CPM ≥ 500

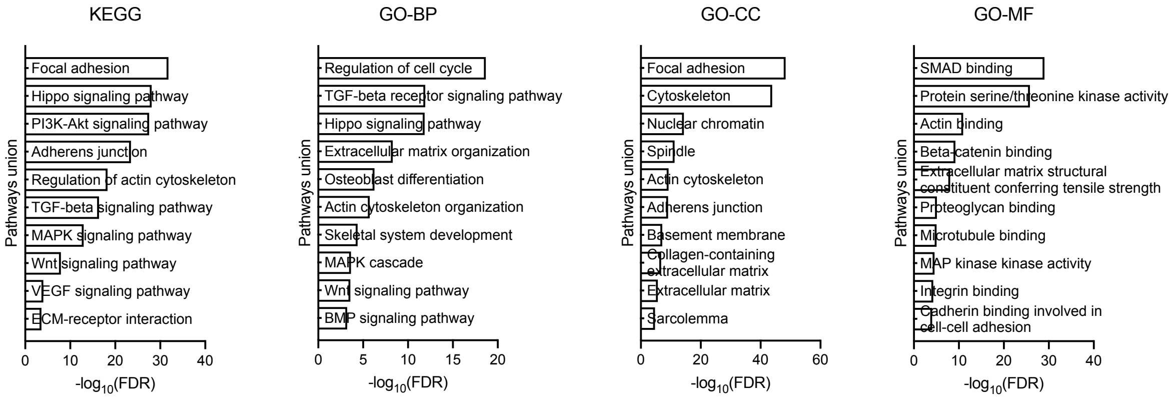


E. 500 > CPM

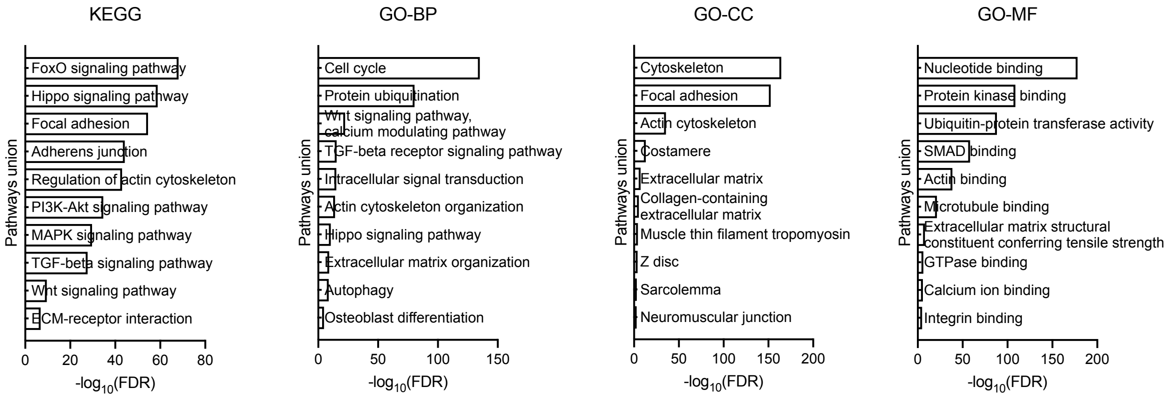


**Figure S3. The top 10 musculoskeletal disease-associated pathways and annotations of the shared miRNAs by abundance.** The KEGG and GO analyses presenting the top 10 musculoskeletal disease-associated pathways and annotations with strong association with the EV cargo miRNAs detected in the Non-deg, Mildly-deg, and Deg samples, as well as the annotation information in the groups with different CPM ranges, **A**, CMP ≥ 25k, **B**, 25k > CMP ≥ 5k, **C**, 5k > CMP ≥ 1k, **D**, 1k > CMP ≥ 500, and **E**, 500 > CMP. GO *p*-value threshold was set at 0.05. The x-axis represents -$\text{Log}_{10}$ (FDR), and the y-axis describes the pathways union of miRNAs involved in the enriched pathways and annotations.

A.

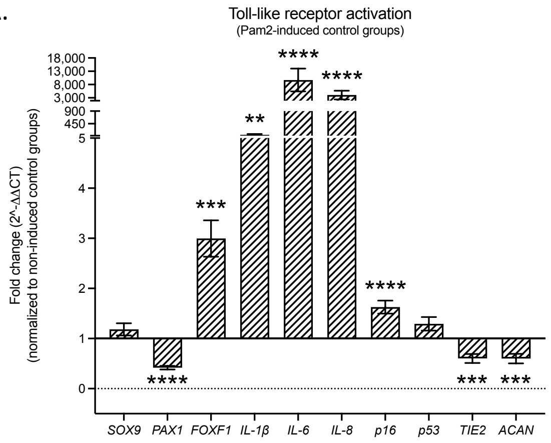


B. Inflammatory mediators and senescent markers (non-inflammatory condition)

a. b. c.

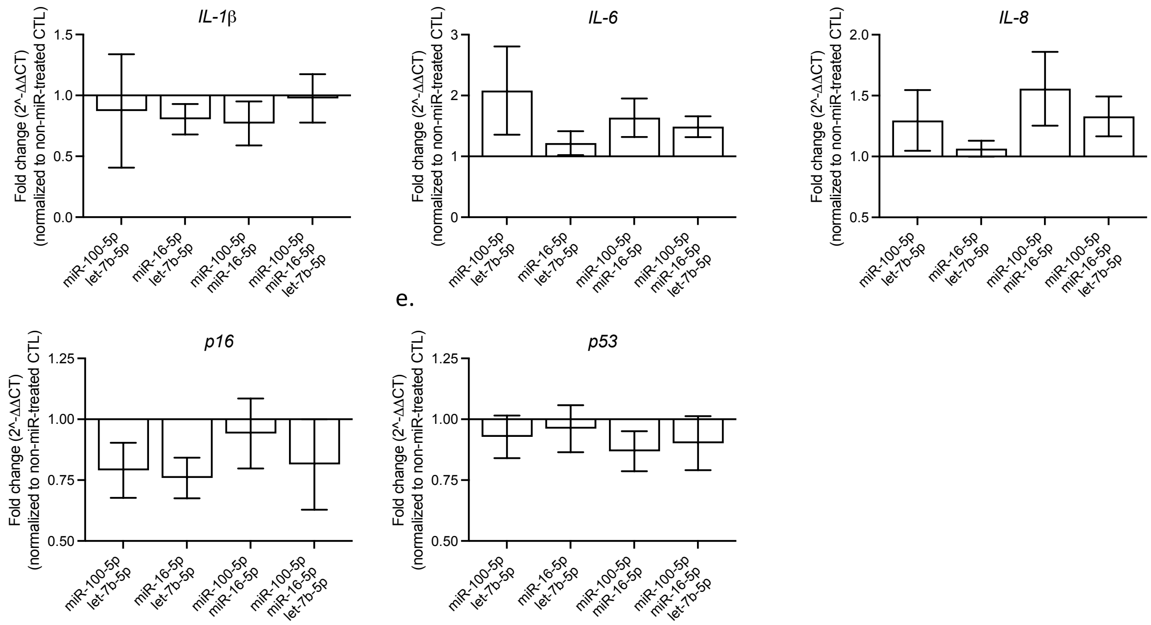


**Figure S4. Toll-like receptor (TLR) activation and the regulatory effects of miRNA mimic combination treatments on inflammatory mediators and senescence markers under non-inflammatory conditions. A.** TLRs were activated by applying 100 ng/mL Pam2CSK4 (Pam2) for 42 hours. The expression of the following genes was compared on human IVD cells with and without TLR activation. Nucleus pulposus (NP) cell phenotypic markers: *SRY-Box Transcription*

*Factor 9 (SOX9)*, *Paired Box 1 (PAX1)*, *Forkhead Box F1 (FOXF1)*, *TEK receptor tyrosine kinase (TIE2)*, and *Aggrecan (ACAN)*. Inflammatory mediators: *Interleukin (IL)-1 beta (IL-1ß)*, *IL-6*, *IL-8*, and senescence markers: *CDKN2A (p16)* and *TP53 (p53)*. **B.** Inflammatory mediator and senescence marker expression, (**a**) *IL-1ß*, (**b**) *IL-6*, (**c**) *IL-8*, (**d**) *p16*, and (**e**) *p53*, of human IVD cells after exposure to miRNA mimic combination treatments in non-inflammatory condition. The patterned and blank histograms represent the inflammatory condition in **A** and non-inflammatory condition in **B**. Data were normalized to cells not exposed to miRNA mimic treatment and are presented as mean ± SEM. *FOXF1* group was assessed by a two-tailed paired Wilcoxon matched-pairs signed rank test and the rest groups were assessed by a two-tailed paired Student's *t*-test in **A.** Data were assessed by a one-way ANOVA with Dunnett's T3 multiple comparisons test in **B**. **, *** and **** indicating a statistical significance of $p < 0.01$, $p < 0.001$, and $p < 0.0001$, respectively. $n = 15$ in **A** and $n = 10$ in **B**.

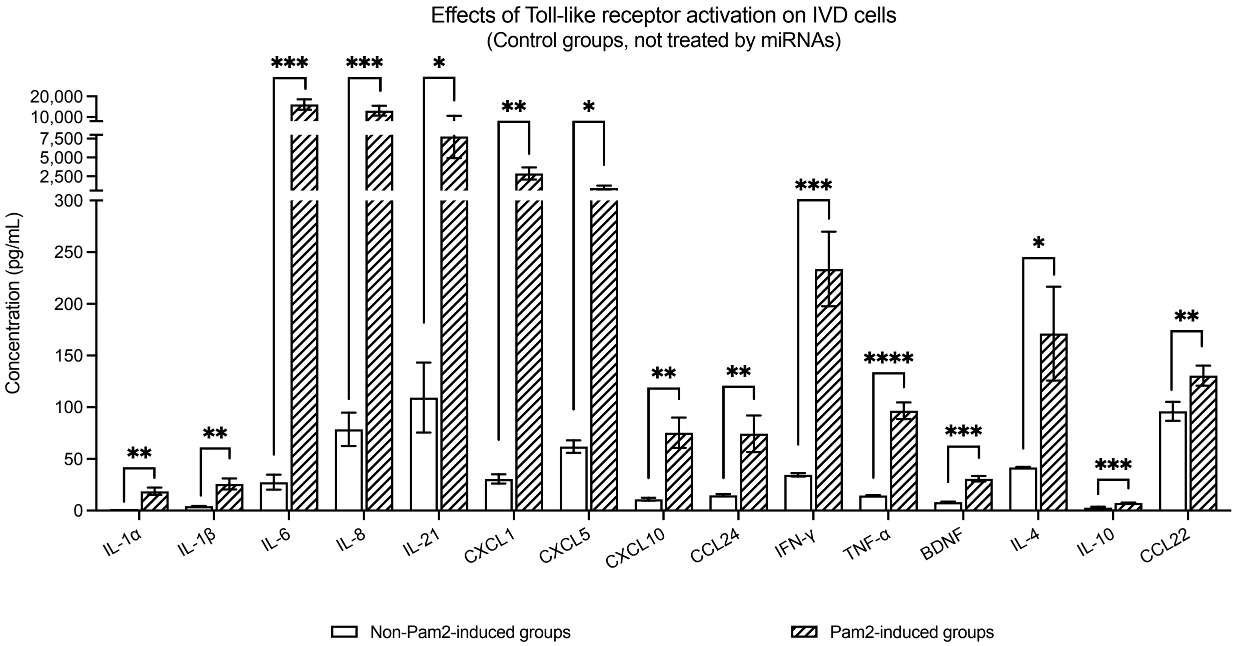


**Figure S5. TLR activation at the protein level.** Comparison of inflammatory mediator production of human IVD cells not exposed to miRNA mimic treatments in non-inflammatory and inflammatory conditions. The blank and patterned histograms represent the non-inflammatory and inflammatory conditions, respectively. Concentration values are presented as mean ± SEM (pg/mL). Data were analysed by a two-tailed paired Student's *t*-test. *, **, ***, and **** indicate a statistical significance of $p < 0.05$, $p < 0.01$, $p < 0.001$, and $p < 0.0001$, respectively. $n = 8$.

**Table S13. Concentration of inflammatory mediators**

| Analyte | Control | Treatment | | | | | |
|---|---|---|---|---|---|---|---|
| | | miR-100-5p | | let-7b-5p | | miR-100-5p + let-7b-5p | |
| | Concentration (pg/mL) | Concentration (pg/mL) | *p*-value | Concentration (pg/mL) | *p*-value | Concentration (pg/mL) | *p*-value |
| **IL-6** | 16,101 ± 2,522 | 5,324 ± 1,223 | 0.0272 | 9,025 ± 1,526 | 0.0416 | 4,324 ± 1,404 | 0.0215 |
| **IL-1α** | 18.67 ± 3.59 | 6.15 ± 1.51 | 0.0053 | 6.11 ± 1.36 | 0.0060 | 4.32 ± 1.20 | 0.0169 |
| **TNF-α** | 96.66 ± 8.09 | 58.01 ± 10.28 | 0.0057 | 61.53 ± 11.29 | 0.0227 | 46.54 ± 11.45 | 0.0240 |
| **IL-1ß** | 25.78 ± 5.25 | 15.59 ± 4.21 | 0.1023 | 17.59 ± 4.76 | 0.1347 | 14.75 ± 4.50 | 0.0438 |
| **IFN-γ** | 233.76 ± 36.13 | 129.75 ± 10.94 | 0.0656 | 140.76 ± 10.84 | 0.1179 | 98.52 ± 35.85 | 0.0253 |
| **IL-10** | 7.47 ± 0.40 | 18.84 ± 3.35 | 0.0351 | 17.19 ± 2.14 | 0.0090 | 17.31 ± 2.81 | 0.0301 |
| **CXCL10** | 75.31 ± 14.62 | 45.14 ± 9.65 | 0.0133 | 42.04 ± 9.83 | 0.0226 | 37.32 ± 8.56 | 0.0103 |
| **IL-8** | 13,100 ± 2,371 | 5,333 ± 2,386 | 0.0778 | 6,325 ± 1,785 | 0.0640 | 3,409 ± 1,569 | 0.0381 |
| **CXCL1** | 2,880 ± 810.10 | 647.48 ± 126.11 | 0.0783 | 682.00 ± 99.04 | 0.0908 | 443.86 ± 113.15 | 0.0497 |
| **CCL24** | 74.33 ± 17.78 | 29.10 ± 4.36 | 0.0691 | 29.22 ± 2.71 | 0.0793 | 22.45 ± 4.36 | 0.0390 |
| **CCL22** | 130.52 ± 9.76 | 427.20 ± 38.04 | 0.0001 | 368.72 ± 24.58 | < 0.0001 | 324.61 ± 33.13 | 0.0015 |
| **BDNF** | 30.81 ± 2.66 | 16.08 ± 2.28 | 0.0219 | 16.59 ± 2.61 | 0.0479 | 11.45 ± 1.83 | 0.0124 |

Concentrations are mean ± SEM.
